# CoLaML: Inferring latent evolutionary modes from heterogeneous gene content

**DOI:** 10.1101/2024.12.02.626417

**Authors:** Shun Yamanouchi, Tsukasa Fukunaga, Wataru Iwasaki

## Abstract

**Motivation:** Estimating the history of gene content evolution provides insights into genome evolution on a macroevolutionary timescale. Previous models did not consider heterogeneity in evolutionary patterns among gene families across different periods and/or clades.

**Results:** We introduce CoLaML (joint inference of gene COntent evolution and its LA-tent modes using Maximum Likelihood), which considers heterogeneity using a Markov-modulated Markov chain. This model assumes that internal states determine evolutionary patterns (i.e., latent evolutionary modes) and attributes heterogeneity to their switchover during the evolutionary timeline. We developed a practical algorithm for model inference and validated its performance through simulations. CoLaML outperformed previous models in fitting empirical datasets and estimated plausible evolutionary histories, capturing heterogeneity among clades and gene families without prior knowledge.

**Availability:** CoLaML is freely available at https://github.com/mtnouchi/colaml.

## 1 Introduction

Tracing the evolutionary histories of gene families is pivotal in phylogenomics. Previous studies have estimated gene gain/loss events and ancestral gene content to address various evolutionary questions, including trait diversity (Thomas *et al*., 2020; Ocaña-Pallarès *et al*., 2022), lifestyle changes (Dharamshi *et al*., 2023) and pathway complexities (Iwasaki and Takagi, 2009). Others have focused more on the origin of life and the gene content of ancient ancestors (Coleman *et al*., 2021; Hyun and Palsson, 2023). The evolutionary predictability of gene content evolution in prokaryotes has been previously assessed (Konno and Iwasaki, 2023). Phylogenetic profiling is another purpose of tracing the evolutionary history of gene families (Pellegrini *et al*., 1999; Moi and Dessimoz, 2023). Functionally related genes often share evolutionary histories; based on this, previous studies have successfully inferred gene functions (Fukunaga and Iwasaki, 2022).

Herein, we focus on the count-based approach, a widely adopted strategy compared with other non-scalable methods, such as phylogenetic reconciliation. In this approach, gene family size (i.e., copy number) or presence/absence is considered using a species tree and ortholog table. Initially, the maximum parsimony (MP) method, a simple method that does not require an evolutionary model, was applied (e.g., Snel *et al*., 2002). Subsequently, model-based methods were introduced to formulate the properties of gene copy number evolution (e.g., Hahn *et al*., 2005). These methods use probabilistic models that incorporate parameters representing features, such as evolutionary rates, typically estimated using the maximum likelihood (ML) criterion.

The evolution of gene families is characterised by heterogeneity in two ways: heterogeneity among genes and over time (heterotachy). First, different genes exhibit different evolutionary patterns: essential genes cannot be lost by definition, whereas accessory ones, such as antibiotic resistance genes, are gained and lost through horizontal gene transfer. Second, the evolutionary rates of gene families vary among lineages and across periods (McCutcheon and Moran, 2012; Inoue *et al*., 2015; Wolf and Koonin, 2013). For instance, the common ancestor of the teleost fish is presumed to have undergone whole-genome duplication (WGD), followed by rapid gene loss (Inoue *et al*., 2015). In addition, symbiotic bacterial clades often undergo extensive genome reduction (McCutcheon and Moran, 2012), and the biphasic model posits that genome evolution alternates between rapid complication and gradual reduction phases (Wolf and Koonin, 2013). Thus, capturing these two types of heterogeneity is key to better understanding gene family evolution.

Existing model-based methods have limited ability to represent such heterogeneities, which involves using multiple gain/loss models or rate categories (Figure 1). For one thing, gene heterogeneity was represented as a mixture model that estimated subpopulations of gene families with different evolutionary rates (Fukunaga and Iwasaki, 2021). However, these cannot account for time heterogeneity since the gain/loss rates are assumed constant across the tree (Figure 1B). For another thing, clade-wise heterogeneity has been addressed by varying the rates at the branch level. Hahn *et al*. (2005) proposed a statistical framework to detect irregular branches by determining whether a branch should be treated separately, implementing it in CAFE. Some models allow for a single branch or a subset of branches to be individually parameterised (e.g., CAFE3 by Han *et al*. 2013 and BadiRate-BR by Librado *et al*. 2012). Other models assign a unique set of parameters to each branch (e.g., Iwasaki and Takagi 2007, BEGFE by Liu *et al*. 2011, and BadiRate-FR by Librado *et al*. 2012; Figure 1C). However, these models do not consider gene heterogeneity. Moreover, specifying different parameters for a subset of branches requires prior knowledge. Conversely, the free rates for each branch often result in too many parameters; therefore, they are not suitable for large datasets. In summary, existing models have difficulty flexibly switching the behaviour of models during evolution while accommodating various gene families.

**Figure 1:**
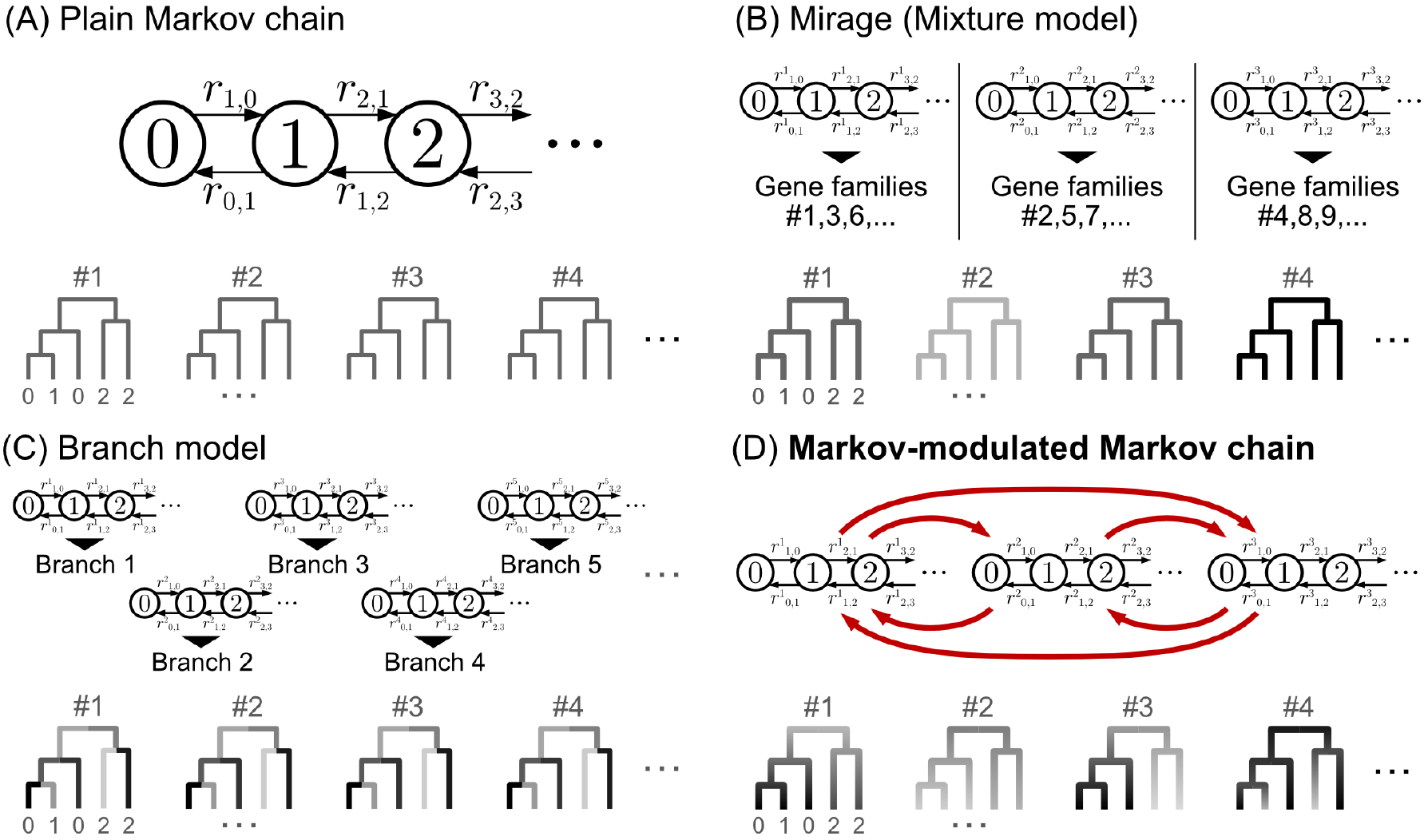
Schematic illustration of gene-content evolutionary models. Enclosed integers and arrows indicate gene copy numbers and state transition events, respectively. The *r* letters on the arrows denote the gene gain/loss rate parameters. Trees display rates assigned to gene families. (A) Plain Markov chain with the most general parameterisation (Kim and Hao, 2014). (B) Mixture model, Mirage (Fukunaga and Iwasaki, 2021). Gene families are clustered into rate categories with no transition in between. (C) Free-rate branch model (Iwasaki and Takagi, 2007; Liu *et al*., 2011; Librado *et al*., 2012). Rate parameters are assigned per branch. (D) The proposed model: Markov-modulated Markov chain. Red arrows indicate category switching.

In this study, we propose CoLaML (joint inference of gene COntent evolution and its LAtent modes using Maximum Likelihood), a probabilistic model that addresses heterogeneity by introducing Markov modulation into gene content evolution modelling. Markov modulation is a statistical technique for modelling model switching that has also been applied to the evolution of sequences and traits (e.g., Baele *et al*., 2021; Boyko and Beaulieu, 2021; Supplementary Notes). This technique enables rate categories (or latent evolutionary modes) to change at any time along tree branches as required for each gene family, thereby representing diverse evolutionary patterns (Figure 1D). We propose an efficient algorithm for model fitting and validate it using simulations. CoLaML showed a better fit to empirical datasets better than previous models, flexibly accommodating both types of heterogeneity.

## 2 Methods

### 2.1 Inputs

The inputs for our model consist of an ortholog table *C*, and a phylogenetic tree *T*. *C* is a *D* × *N* integer (ordinal data) matrix, where *D* and *N* denote the numbers of species (genomes) and gene families (orthogroups), respectively. The (*d, n*)-th element of *C* represents the observed copy number of *n*-th gene family in *d*-th genome. We assume that 0 ≤ *C*_*d,n*_ ≤ *l*_max_ for all (*d, n*), such that there are *L* := *l*_max_ + 1 possible values at most. To satisfy this requirement, the values exceeding *l*_max_ are converted to *l*_max_.

*T* is a rooted tree with a fixed topology, *D* leaves, and *M* − *D* internal nodes. Let *t*_*m*_ be the branch length between the *m*-th node and its parent; *T* can be non-binary (i.e., polytomy is allowed). Our model relies on a nonstationary Markov chain and is sensitive to the rooting of *T*. The choice of branch lengths *{t*_*m*_*}* substantially affects the results. We recommend rescaling them using, for example, changes averaged over the maximum parsimonious reconstructions (see Supplementary Materials §3.5).

### 2.3 Probabilistic model of gene copy number evolution

We began by modelling changes in gene copy numbers in each infinitesimal time step. Typically, previous models have been based on continuous-time Markov chains (CTMCs), where states and their transitions represent gene copy numbers and their changes, respectively. The rates of these changes are specified by a transition rate matrix *R* of size *L*× *L*: the off-diagonal (*i, j*)-th element of *R* represents the per-time flow of state probability from *i* to *j*, whereas the *j*-th diagonal element, defined as −∑ _*i*:*i* ≠*j*_ *R*_*i,j*_, represents the outflow of state probability *j*. Since the gene copy number is an ordinal quantity, *R* is a tridiagonal matrix that restricts instantaneous transitions to adjacent copies.

Using Markov modulation, we extended CTMC to incorporate heterogeneity. Markov modulation allows CTMC to have different states even with the same copy number. Specifically, our model has multiple (hereafter *K*, which is user-defined) states for each copy number, constituting *K* rate categories that represent the latent evolutionary modes. Gene gain/loss within each rate category is modelled using plain CTMCs as in previous models, such as the birth-and-death process with all rates different (BDARD; Kim and Hao, 2014). Furthermore, Markov modulation allows for transitions among *K* states of the same copy number, which enables model switching during evolution (Figure 1D). Accordingly, we obtain a transition rate matrix of the Markov-modulated CTMC:

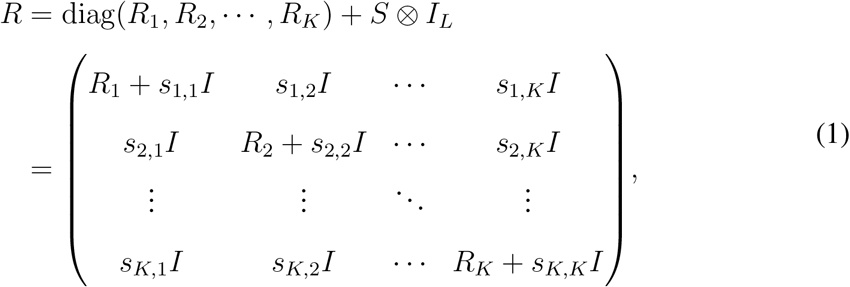

where ⊗ is the Kronecker product, *R*_1_, *R*_2_, · · ·, *R*_*K*_ are *L* × *L* transition rate matrices of the plain CTMC for copy number evolution, and *S* is a *K* × *K* transition rate matrix for model switching. The elements are denoted by *r*_*k*;·,·_ and *s*_·,·_, satisfying ∑_*i*_ *r*_*k*;*i,j*_ = 0 for all *k* and *j* and ∑_*k*_ *s*_*k,l*_ = 0 for all *l*. Hereafter, we use four subscripts for the elements of *R*: *R*_*k,i*;*k,j*_ = *r*_*k*;*i,j*_ (×*;i* − *j*×; = 1) and *R*_*k,i*;*l,i*_ = *s*_*k,l*_. Herein, *R* is left unrestricted in terms of gene gain/loss and model switching to consider the most general case.

Next, we extended the Markov-modulated CTMC to a tree. First, we considered the evolution of a single gene family on a single branch. Given the initial copy number *y*, initial category *b*, and rate matrix *R*, the system follows the differential equation *dP* (*t*)*/dt* = *RP* (*t*), where *P* (*t*), formally *P* (*x, a* ×; *y, b, tR*), is the probability of state (*x, a*) at time *t*. Since (*x, a*) =(*y, b*) at *t* = 0, we obtain *P* (*x, a* ×; *y, b, tR*) =[exp(*tR*)]_*x,a*;*y,b*_. The likelihood of each gene family can then be obtained by enumerating all possible combinations of states at the tree nodes and summing the products of the transition probabilities exp(*t*_*m*_*R*) weighted by the initial (root) state probability *πϕ*, where *ϕ*_*y*_ is the probability of *y* and *π*_*y*;*b*_ is the probability of *b* given *y*. This can be calculated using the so-called inside algorithm, as described by Kiryu (2011), with slight modifications (see Supplementary Materials for further details). Finally, assuming that all gene families are independent and identically modelled, the likelihood over the dataset is the product of the likelihood of each gene family.

### 2.3 Parameter estimation with EM algorithm

Next, we estimated the model parameters *θ* := (*r*_*k*;*i,j*_, *s*_*k,l*_, *π*_*x*;*a*_, *ϕ*_*x*_) using the maximum likelihood (ML) method.

Let us reinterpret the marginal likelihood described above as the sum (integral) of probabilities over all paths. Let *X* be a path (i.e., evolutionary history) along *T* that is consistent with *C*_*n*_, the *n*-th column of *C*. The probability of *X* is given by

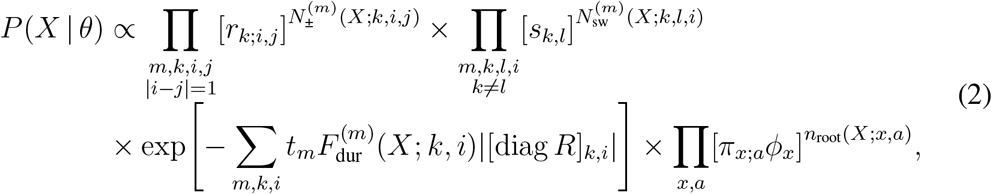

where *N*_±_, *N*_sw_, *F*_dur_, and *n*_root_ constitute sufficient statistics for *X*. 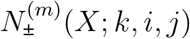 and 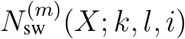 represent the number of state transitions (*k, j*) → (*k, i*) (copy number increase/decrease) and (*l, i*) → (*k, i*) (category switching) on branch *m*, respectively. 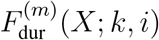 represents the total proportion of time for which state (*k, i*) lasts along branch *m. n*_root_(*X*; *x, a*) is the indicator that returns 1 if the state of *X* is (*x, a*) at the tree root, and 0 otherwise.

In effect, we cannot observe *X* directly, except for its extant copy number states *C*_*n*_; nevertheless, we can estimate the parameters using the expectation-maximisation (EM) algorithm (Dempster *et al*., 1977). This algorithm is often used to obtain ML estimates of parameters based on data containing unobservable values, as in our case. To refine the parameters iteratively, two steps are alternately repeated: expectation (E) and maximisation (M).

The E-step computes the expected value of the sufficient statistics conditioned by *C*_*n*_:

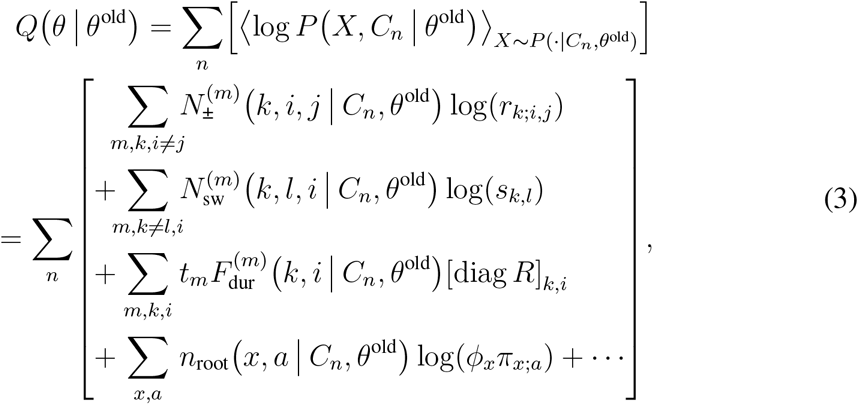

where [*N*×*;F*×*;n*](· · · ×; *C*_*n*_, *θ*^old^) represents the expected value of the sufficient statistics in Equation (2). These expected values are computed efficiently (Kiryu, 2011) as detailed in Supplementary Materials. Meanwhile, in the M-step, the parameters are updated to maximise the *Q*-function, as described in Supplementary Materials.

Furthermore, we evaluated the time complexity of the proposed algorithm. The most time-consuming step was the computation of the expected number of transitions. In our model, each node has *KL* possible states, and instantaneous transitions can occur in 2*K*(*L* − 1) ways for gene gain/loss and *K*(*K* − 1)*L* ways for category switching. Considering all the combinations (i.e., node states at both ends of a branch and instantaneous transitions for all branches and gene families), our model has a time complexity of *O*(*NMK*^4^*L*^3^) per iteration for the EM algorithm.

### 2.4 Reconstruction of evolutionary history

The ancestral states were estimated using two classical methods with the estimated parameters. Joint reconstruction identifies the most probable course of evolution to estimate the unobserved states at the tree nodes. Efficient algorithms are described in similar settings (e.g., Pupko *et al*., 2000; Fukunaga and Iwasaki, 2021) and can be easily extended to our case. Marginal reconstruction estimates the most probable state individually for each (internal) node by marginalising all unobserved states except their own. This is achieved by determining the state that maximises the posterior state probability for each node.

### 2.5 Simulation datasets

The performance of CoLaML was evaluated using synthetic ortholog tables and their underlying evolutionary histories obtained from simulations. The simulations consisted of 19 conditions designed to assess the individual effects of four factors: *l*_max_ (1–6), *K* (1–6), *D* (32–512; 2x steps), and *N* (1,000–16,000; 2x steps), with (*l*_max_, *K, D, N*) = (2, 3, 128, 4000) as the baseline.

Ten datasets were generated for each condition using random trees and parameter sets. Phylogenetic trees were generated via the Yule process (birth rate 1) using the rphylo function in ape (Paradis and Schliep, 2019) in R. Rate parameters *r* and *s* were sampled from the gamma distributions Γ(2, 0.5) and Γ(3, 0.1*/*3), respectively. The initial category probability *ϕ* was drawn from a symmetric Dirichlet distribution Dir([20, · · ·, 20]). The Initial copy number probabilities *π* were sampled from a Dirichlet distribution with ad hoc concentration parameters derived by discretising a log-normal distribution (scale 0.8, shift 0.5): 20 [.193, .501, .180, .067, .029, .014, .017] when *l*_max_ = 6; otherwise, extra parameters were deleted and added to the final (i.e., *l*_max_-th) value.

Model fitting was performed as outlined in Supplementary Materials. Briefly, for each condition, we conducted 10 EM runs and selected the run with the highest final likelihood. Aborted runs that occurred occasionally owing to the numerical instability of eigendecomposition, possibly due to unrealistically low rate parameters, were excluded from the analysis.

Our model is arbitrary in terms of category permutation. To handle label switching, we mapped categories between the simulation (as ground truth) and estimation results by solving the linear sum assignment problem to minimise the total absolute percentage error of the gain/loss rates ∑ ×*;r*_est._ − *r*_truth_×*;/*×*;r*_truth_×;.

The parameter estimation accuracy was assessed using the mean absolute percentage error (MAPE) for rates *r* and *s* and the Earth mover’s distance (EMD) for root probabilities *π* and *ϕ*. For *π*, the unit transfer cost from copy number *a* to *b* was defined as the Manhattan distance *d*(*a, b*) = ×*;a*− *b*×;. For *ϕ*, the unit transfer cost from category *x* to *y* was defined as the discrete metric *d*(*x, y*) = [*x* ≠ *y*], where [·] is the Iverson bracket (i.e., the similarity between rate categories was not considered).

To evaluate execution time, we measured the CPU time required for 50 EM iterations using the same series of simulations with 19 conditions × 10 replicates. The measurements were performed in a random order and repeated 10 times. We suppressed the multithreading and precompilation by dependent packages, such as numpy, scipy and numba, to obtain the net time in a single thread. The computations were performed on a server with four Intel^®^ Xeon^®^ CPU E5-4627 v3, with the CPU affinity set to run on a single NUMA node. The results reported herein are based on CoLaML v0.1.dev14+g6c01617 on Python v3.10.

### 2.6 Empirical datasets

We compiled two empirical datasets, fish and bacteria. The fish dataset was constructed from the Fish Tree of Life (Rabosky *et al*., 2018) and OrthoDB v11 (Kuznetsov *et al*., 2023). By cross-referencing Actinopterygii (ray-finned fish) species names in OrthoDB with the Fish Tree of Life, we identified exact matches or synonyms for 128 out of 135 species, mapping four to congeneric species and discarding three. We then checked in NCBI RefSeq for the latest status (as of 6 July 2024) of the OrthoDB source assemblies. We removed genomes tagged as “contaminated”, except for a basal species *Acipenser ruthenus* (sterlet) due to its lineage-specific WGD event. We also excluded genomes from Euteleostei with “Contig” or “Scaffold” assembly status for down-sampling. Finally, we retrieved Actinopterygii-level orthologous groups from OrthoDB for the remaining 104 species to obtain their copy numbers.

The bacteria dataset was generated based on the results of Coleman *et al*. (2021). We downloaded the Extended Data from the Figshare repository and extracted copy number profiles of Clusters of Orthologous Genes (COGs) from reconciled gene trees. We adopted the phylogeny placing *Fusobacterium* as the sister group of *Gracilicutes* (i.e., “Root 1” in their main article, or the rooting by node “528” in their supplements). For the COG categories, we relied on eggNOG annotations rather than NCBI COGs, as there were a few inconsistencies.

Gene families in <5% of the subject species were excluded from both datasets to reduce noise from taxonomically restricted genes. The resulting datasets comprised 104 species × 25,530 families in the fish dataset and 265 species × 2,876 families in the bacteria dataset.

We pre-processed the input phylogenetic trees and adjusted the branch lengths according to evolutionary changes in copy numbers approximated by the maximum parsimony method. For further details see Supplementary Materials (§3.5; Supplementary Figures S9, S11, and S12).

We performed five-fold cross-validation to compare the performance of the evolutionary models. First, we randomly divided the gene families in the empirical datasets into five subsamples of equal size. Next, we applied the EM algorithm using the four subsamples as training data. Finally, we tested the models using the estimated parameters by calculating the likelihood of the remaining subsample. We repeated this process five times, ensuring that each subsample served as test data. Overall goodness-of-fit scores were obtained as the sum of the (total) log-likelihoods for each fold.

## 3 Results

### 3.1 Performance of CoLaML evaluated by simulations

We validated our model using simulation analysis. Supplementary Figure S1 shows the accuracy of the parameter estimation under several settings. Errors in the rate parameters *r* and *s* were measured using the mean absolute percentage error (MAPE), whereas those in the root probabilities *π* and *ϕ* were measured using the Earth mover’s distance (EMD). EMD is a dissimilarity measure between two distributions defined as the minimal cost of transporting a mass to match one distribution to another. In our cost setting, EMD takes values in the range of [0, *l*_max_] for *π* and [0, 1] for *ϕ*.

In the baseline setting (*l*_max_, *K, D, N*) = (2, 3, 128, 4000), the median parameter estimation errors were 0.031 for *r*, 0.60 for *s*, 0.050 for *π* and 0.024 for *ϕ*, respectively. The lower accuracy for *s* compared with *r* was presumably due to fewer transition events and the unobservability of the rate categories.

Our model exhibited good performance in the middle range of *l*_max_ (2–4; Supplementary Figure S1A). The accuracy deterioration for *r* at larger *l*_max_ (e.g., 0.084 when *l*_max_ = 6) may be attributed to fewer transition events at high copy numbers in our simulations (see Methods). The poor estimation at *l*_max_ = 1 (*r*: 0.066, *s*: 2.1, *π*: 0.036, and *ϕ*: 0.073) is likely due to identifiability issues arising from the data generation procedure, and not from the model itself. The random model parameters in the simulations may result in multiple rate categories with similar gain and loss rates.

As expected, a larger *K* reduced the estimation accuracy (e.g., *r*: 0.14, *s*: 1.5, *π*: 0.12, and *ϕ*: 0.067 when *K* = 6; Supplementary Figure S1B), likely due to the increased model complexity, especially in category switching.

Larger *D* and *N* values resulted in better parameter estimation (Supplementary Figure S1C and D). Smaller *D* led to more severe accuracy deterioration (e.g., *r*: 0.25, *s*: 2.6, *π*: 0.14 and *ϕ*: 0.13 when *N* = 32) than smaller *N* (e.g., *r*: 0.073, *s*: 0.92, *π*: 0.11 and *ϕ*: 0.050 when *D* = 1000). This indicates the importance of the total branch length and/or number of branches (i.e., the number and/or independence of evolutionary events) for parameter estimation. Overall, these results suggest that the proposed method accurately estimates the model parameters, except under unrealistic and extreme conditions.

Supplementary Figures S2–S4 illustrate the fraction of correctly estimated ancestral states under various conditions using either joint or marginal reconstruction. Copy numbers were accurately estimated, with a median accuracy >80% in all conditions, even at root nodes. However, the accuracy decreased from leaves to the root as it became distant from observable data. The accuracy of copy number estimation remained largely unaffected by hyperparameters nor reconstruction methods.

In contrast, the rate category estimation was not less accurate and exhibited considerable variation depending on the conditions and replicates. Even at the baseline (*l*_max_, *K, D, N*) = (2, 3, 128, 4000), the accuracy ranged from approximately 50 to 90%.

The rate category estimation was strongly influenced by *K* (larger corresponding to harder) and *D* (larger corresponding to better), whereas *l*_max_ (hard when small) and *N* (larger corresponding to better) had milder impacts.

The difficulty in estimating the rate categories likely arose from their inherent unobservability, which required an indirect inference. In model-based (ancestral) state estimations, errors in the parameter estimation propagate to the evolutionary history estimation. Moreover, unlike copy numbers, rate categories cannot be observed even at leaf nodes, making their estimation more unstable.

Regarding the reconstruction methods, marginal reconstruction outperformed joint reconstruction, even under challenging conditions. In principle, joint reconstruction explores a representative path in evolutionary history, whereas marginal reconstruction averages all possible paths. The latter strategy appeared to be more effective, at least in estimating the states independently per node.

Finally, we evaluated the computational time required for CoLaML (Supplementary Figure S5). The required time was nearly linear for *D* and *N*, consistent with the theoretical complexity. We confirmed that the execution time was more than linear for *l*_max_ and *K*. Although the time complexity should theoretically be proportional to the third power of *l*_max_ and the fourth power of *K*, these factors had weaker influences than expected. This suggests that when *l*_max_ and *K* are not very large the effect of steps irrelevant to the asymptotic bottleneck (e.g., the inside-outside algorithm) is not negligible.

### 3.2 Fit to empirical data compared with existing models

Next, we assessed the ability of CoLaML to represent real-world data by comparing CoLaML with our previous models (Iwasaki and Takagi, 2007; Fukunaga and Iwasaki, 2021) through likelihood-based cross-validation using empirical datasets.

Figure 2 shows the scores of the models for various settings. In all settings, the plain CTMC, a special case of Mirage (the mixture model; Fukunaga and Iwasaki, 2021) and CoLaML with a single rate category, had the lowest score and degree of freedom (dof). The free-rate branch model (Iwasaki and Takagi, 2007) scored second lowest despite its large dof, with the score not even available for the bacteria dataset when *l*_max_ = 3 due to numerical instability. This suggests that the branch model overfits and is unsuitable for datasets with a large *D*. Mirage and CoLaML scored higher as the number of rate categories increased (i.e., as they become more flexible). At a similar complexity, CoLaML consistently outperformed Mirage, indicating that our model better captured the evolutionary patterns in the empirical datasets.

**Figure 2:**
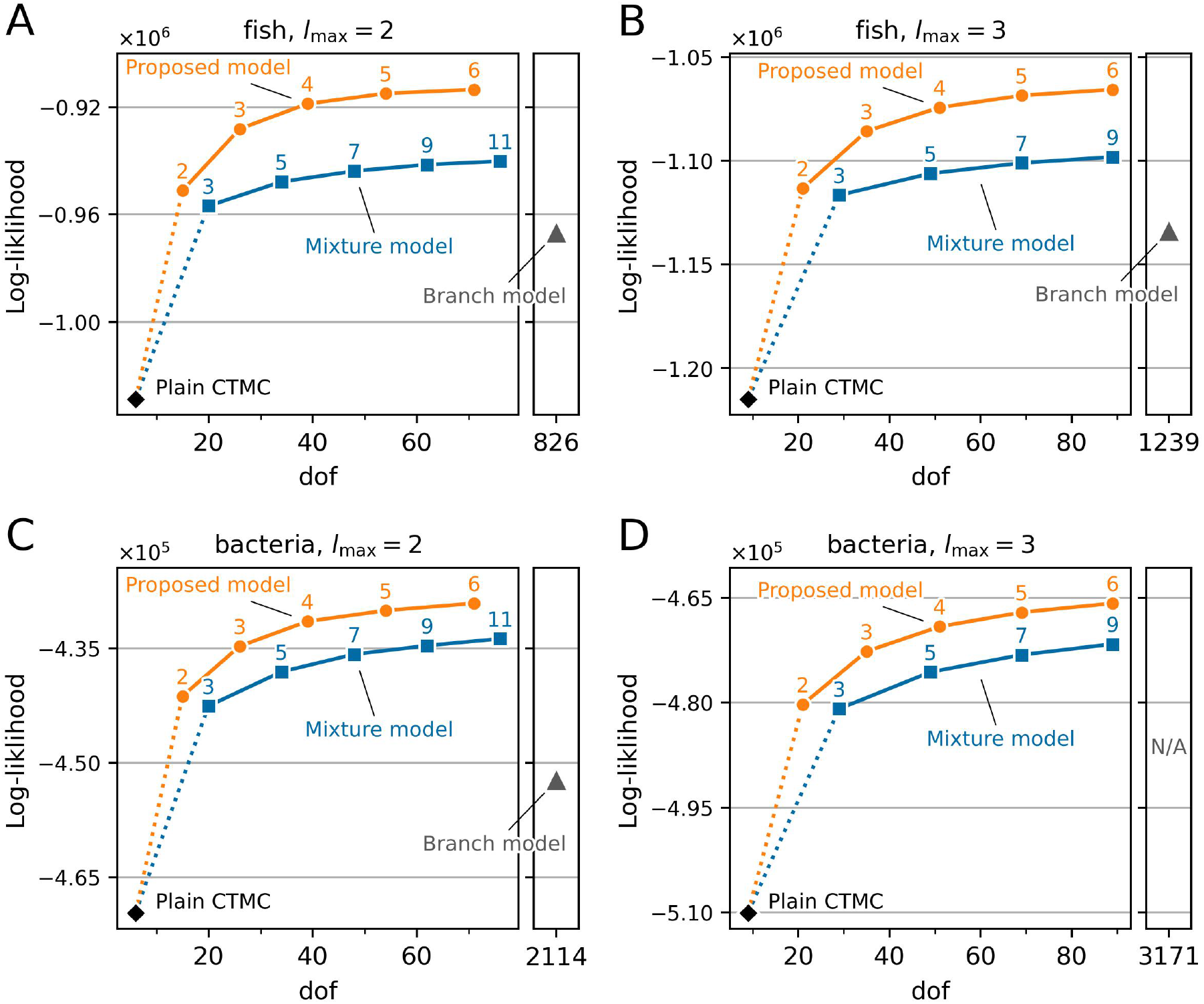
Model fit comparison through cross-validation. The x-and y-axes represent the degree of freedom (dof) of the models and the goodness-of-fit score measured by likelihood-based cross-validation, respectively. The numbers next to the plots indicate the number of rate categories (CoLaML) or mixtures (Mirage; Fukunaga and Iwasaki, 2021).

### 3.3 Estimated ancestral gene content and rate categories

We further investigated how our model captured heterogeneity in the empirical datasets. Figure 3A shows the estimated parameters for the fish dataset with *l*_max_ = 2 and *K* = 3 as this setting provided reasonable computational time and biological interpretation. Three categories can be interpreted as #1: the “fast-evolving” mode with rapid changes in copy numbers, #2: the “single-copy” mode with frequent transition from any to 1 but not in reverse, and #3: the “duplication” mode characterised by rapid transition from 1 to 2. In line with this interpretation, massive transitions from categories #1 to #2 were observed (Figure 3C) alongside lineage-specific WGD events (Figure 3B; 3R(s) in sturgeon/paddlefish, 4R in carps, and 4R in salmonids). Additionally, a slight increase in the frequency of category #2 was observed in the branch leading to the last common ancestor of teleosts. This appears to reflect a trace of 3R in teleosts (i.e., genes not lost soon after) obscured by deep branching. These results demonstrate that CoLaML successfully captures varying evolutionary patterns across lineages.

**Figure 3:**
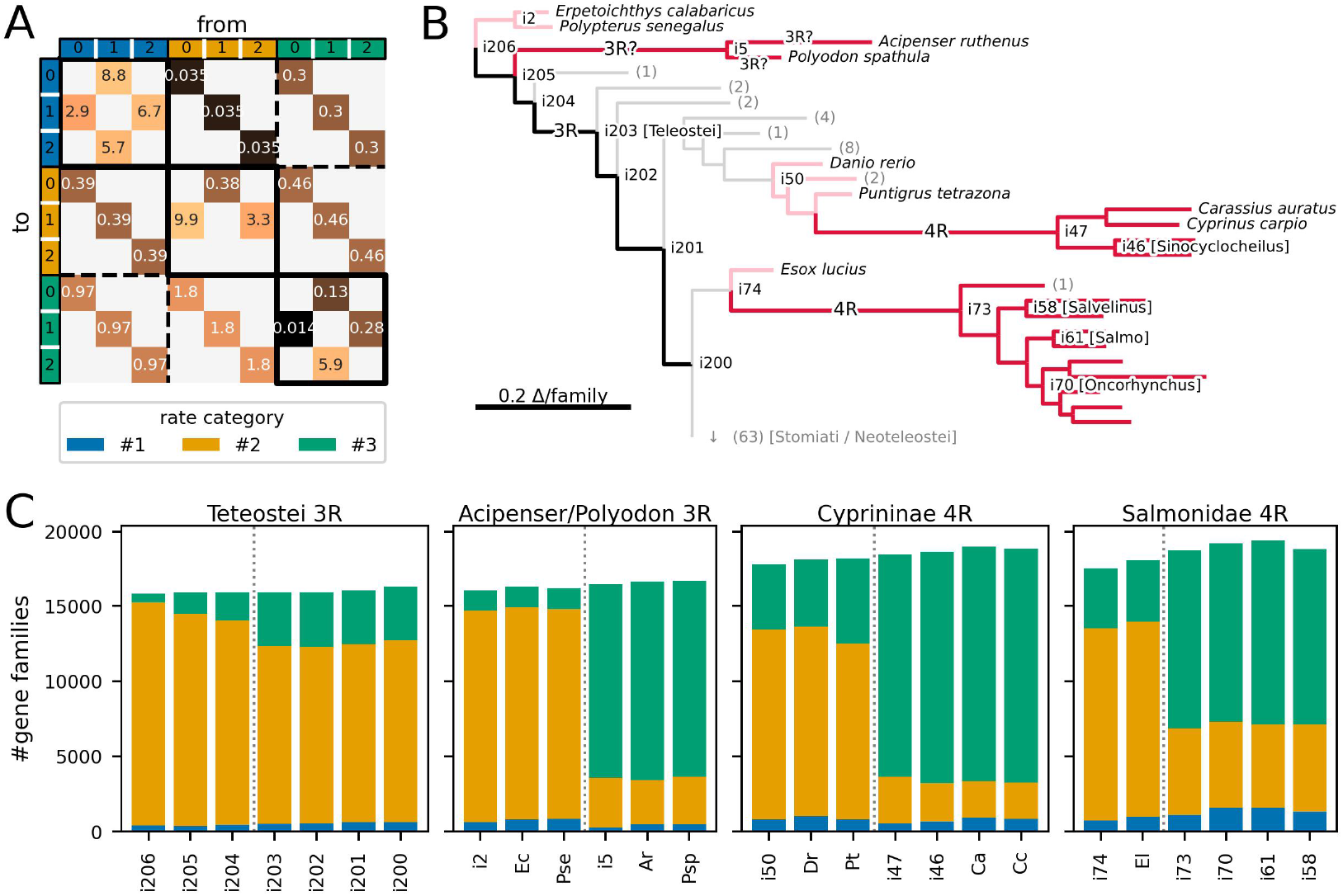
Application of CoLaML to the ray-finned fish dataset. (A) Maximum likelihood estimation of transition rate matrix *R*. (B) Simplified phylogram of species with associated whole-genome duplication (WGD) events (“3R”s and “4R”s). Clades irrelevant to WGDs are collapsed, yielding the number of species therein. Branch lengths are scaled to mean parsimonious changes in copy number per gene family. Clades and paths of particular relevance to WGD events are highlighted. (C) Distributions of estimated rate categories of gene families before and after WGD events. Node labels are based on (B), with extant species abbreviated by their initials. Bar heights represent the number of gene families in the genome at each node.

Figure 4A shows the estimated parameters for the bacteria dataset with *l*_max_ = 2 and *K* = 3. Category #1 exhibited almost no transition between 0 and 1 but rapid transitions from 2 to 1 (the “near-static” mode at 0 or 1 copy). Categories #2 and #3 had similarly high transition rates between 0 and 1 but differed in the rates between 1 and 2 (the “dynamic” modes with/without duplication).

**Figure 4:**
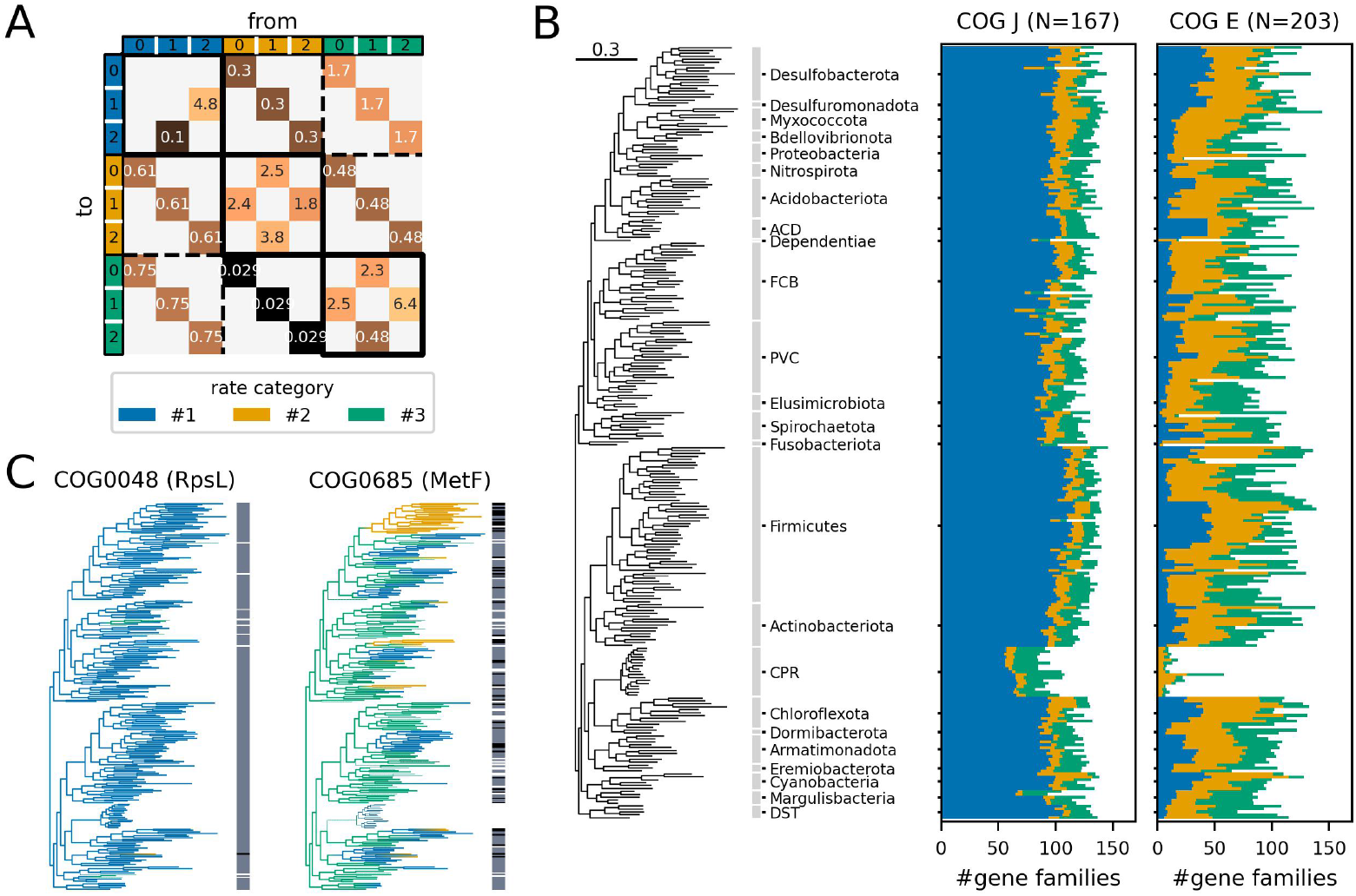
Application of CoLaML to the bacteria dataset. (A) Maximum likelihood estimation of the transition rate matrix *R*. (B) Distributions of estimated rate categories for gene families of COG categories J (Translation, ribosomal structure and biogenesis) and E (Amino acid transport and metabolism) in each extant species’ genome, ordered by phylogenetic relationships based on Coleman *et al*. (2021). Bar widths represent the number of gene families present in each genome. (C) Examples of evolutionary history estimation of rate categories. Branches are coloured by the most likely categories of nodes at both ends, drawn in a thinner line where the copy number is estimated to be 0. Bands alongside the tree indicate current copy numbers (white for 0, grey for 1 and black for 2).

A notable difference between these results and those from the fish dataset was the absence of a genome-scale shift in the rate categories (Supplementary Figure S11). This may be surprising at first sight since several bacterial clades are known for substantial genome reduction (i.e., gene-loss evolutionary mode). Instead, we found that gene families in different functional categories tended to show different evolutionary patterns (Figure 4B). In other words, among-gene heterogeneity was prioritised over clade-wise heterogeneity in this dataset. The copy numbers of essential genes remains unchanged even in reduced genomes, such as those of the candidate phyla radiation (e.g., COG0048; RpsL: 30S ribosomal subunit protein S12), whereas non-essential genes are characterised by evolutionary modes that are dependent on phylogenetic clades (e.g., COG0685; MetF: 5,10-methylenetetrahydrofolate reductase) (Figure 4C). It should be noted that such essential differences in the patterns of gene copy number evolution could not be revealed without CoLaML.

## 4 Discussion

In this study, we developed CoLaML, an unexplored class of gene content evolutionary models incorporating Markov modulation, that addresses heterogeneity by switching between the latent evolutionary modes that govern evolutionary patterns. Simulations demonstrated that the parameters and ancestral states could be accurately estimated within a feasible time. Analyses with empirical datasets showed that our model could recognise the inherent heterogeneity among genes and lineages in an unsupervised manner.

Our method can be extended to provide confidence intervals, whereas the proposed model only provides point estimates. Confidence intervals are particularly helpful for category switch rates as their estimation accuracy is relatively limited (Supplementary Figure S1). Potential approaches include resampling (e.g., bootstrapping) and Bayesian methods. Future studies should focus on improving the computational efficiency of resampling and exploring suitable Bayesian priors.

Determining the number of rate categories *K* in our model is a non-trivial task. Our results showed that the likelihood was not saturated in the cross-validation within the explored range (Figure 2), suggesting that the datasets are better represented by more complex models with a larger *K* at least in terms of likelihood. However, such complex models would incur considerable computational costs and compromise interpretability. Thus, the balance between these factors remains to be considered.

Another concern is the identifiability issue of the proposed model. In addition to the trivial cases of label switching, it remains to be confirmed whether entirely different parameterisation (i.e., different configurations of rate categories) can equivalently explain the observations. Ensuring a model’s unique interpretation is crucial; otherwise, the rate categories obtained may not be interpreted as biologically meaningful evolutionary modes. General theories of the identifiability of Markov-modulated Markov chains (Ito *et al*., 1992; Larget, 1998) may help to verify this mathematically.

One potential application of our model is phylogenetic profiling. Indeed, CoLaML better represented the empirical datasets than the other models (Figure 2), distinguishing evolutionary patterns between gene families of different functions (Figure 4). Therefore, the evolutionary histories estimated by our model can potentially better predict the functional relationships among gene families.

## Acknowledgements

The authors thank Cosentino Salvatore for comments to codes. We would like to thank Editage (www.editage.jp) for English language editing.

## Software and data availability

The Python implementation of CoLaML is freely available at https://github.com/mtnouchi/colaml. Datasets and analysis codes are available at https://github.com/mtnouchi/colaml-test.

