## Supplementary Materials for "CoLaML: Inferring latent evolutionary modes from heterogeneous gene content"

Table 1: Quick reference for notation in this article

| Symbol | Description |
|---|---|
| $T$ | Phylogenetic tree |
| $C$ | Ortholog table |
| $D$ | #(leaves in $T$) = #(rows in $C$) |
| $M$ | #(nodes in $T$) |
| $N$ | #(gene families) = #(columns in $C$) |
| $l_{\max}$ | Max gene copy number |
| $K$ | #(rate categories) |
| $m$ | Node/Branch index for $T$ |
| $\mathrm{pa}(m)$ | Parent node of node $m$ |
| $\mathrm{ch}(m)$ | Set of child nodes of node $m$ |
| $\mathrm{sib}(m)$ | Set of sibling nodes of node $m$ (i.e., $\mathrm{ch}(\mathrm{pa}(m)) \setminus \{m\}$) |
| $\mathrm{sub}(m)$ | Set of nodes in the subtree rooted by $m$ |
| $t_m$ | Branch length between nodes $m$ and $\mathrm{pa}(m)$ |
| $n$ | Gene family index |
| $C_n$ | Copy-number profile of gene family $n$ (i.e., $n$-th column of $C$) |
| $\mathrm{inside}_m(C_n)$ | Partial $C_n$ restricted to descendant leaves of $m$ |
| $\mathrm{outside}_m(C_n)$ | Complement of $\mathrm{inside}_m(C_n)$ |
| $i, j/a, b$ | Copy number index (instantaneous/tree-nodal) |
| $k, l/x, y$ | Category index (instantaneous/tree-nodal) |
| $r_{k;i,j}$ | Gain/Loss rate ($j \to i$; category $k$) |
| $s_{k,l}$ | Category switching rate ($l \to k$) |
| $\pi_{x,a}$ | Probability of copy number $a$ at the root in category $x$ |
| $\phi_x$ | Probability of category $x$ at the root |
| $\theta$ | All parameters to be estimated $(r, s, \pi, \phi)$ |
| $F_{\mathrm{dur}}$ | Fraction of each state's total duration (holding time) |
| $N_{\pm}$ | Number of state transition for gene gain/loss |
| $N_{\mathrm{sw}}$ | Number of state transition for category switching |

### 1 Supplementary Methods

#### 1.1 Inside-Outside algorithm

##### 1.1.1 Description

Let $m$ be the node index of input tree $T$, $\mathrm{pa}(m)$ be the parent of $m$, and $\mathrm{ch}(m)$ be the set of child nodes of $m$. We denote the branch length between $m$ and $\mathrm{pa}(m)$ as $t_m$.

First, we define the inside variables $\alpha$ as:

$$\alpha_n(m, x, a) := P\big(\mathrm{inside}_m(C_n) \,\big|\, \text{state at } m = (x, a), \theta\big), \tag{S1}$$

where $\mathrm{inside}_m(C_n)$ denotes the partial observations of $C_n$ from the species corresponding to the descendant leaf nodes of $m$. By definition, if $m$ is a leaf node, $\alpha_n(m, x, a) = [C_{m,n} = a]$, otherwise, $\alpha_n$ satisfies the following recursive relationship:

$$\alpha_n(m, y, b) = \prod_{m' \in \mathrm{ch}(m)} \sum_{x,a} \alpha_n(m', x, a) \cdot P(x, a \,|\, y, b, t_{m'} R). \tag{S2}$$

Therefore, we can recursively obtain the inside variables for all tree nodes, starting from the tree leaves and proceeding towards the root (i.e., by post-order traversal).

The characteristic point of our model is the presence of more than one—more specifically, $K$—nonzero inside variables at tree leaves for each $n$. Nonetheless, the inside algorithm is feasible in the same manner as in ordinary CTMCs.

Next, we define the outside variable $\beta$ as follows:

$$\beta_n(m, x, a) := P\big(\mathrm{outside}_m(C_n), \text{state at } m = (x, a) \,\big|\, \theta\big), \tag{S3}$$

where $\mathrm{outside}_m(C_n)$ denotes the partial observations of $C_n$ from the species corresponding to the leaf nodes not descending from $m$. Consequently, $\beta_n$ satisfies the following recursive relationship:

$$\begin{aligned}\beta_n(m, x, a) = \sum_{y,b} \Bigg[ & P(x, a \,|\, y, b, t_m R) \cdot \beta_n(\mathrm{pa}(m), y, b) \\ & \times \prod_{m' \in \mathrm{sib}(m)} \sum_{x',a'} \alpha_n(m', x', a') \cdot P(x', a' \,|\, y, b, t_{m'} R) \Bigg],\end{aligned} \tag{S4}$$

where $\mathrm{sib}(m) := \mathrm{ch}(\mathrm{pa}(m)) \setminus \{m\}$ represents the sibling node set of the node $m$. Accordingly, the outside variables can be calculated recursively for all tree nodes from the root ($\beta_n(\mathrm{root}, x, a) = \pi_{x;a}\phi_x$) to the leaves (i.e., by pre-order traversal).

Using the inside and outside variables, we can calculate several important probabilities such as

- the likelihood of column $C_n$:

$$P(C_n \mid \theta) = \sum_{x,a} \alpha_n(\mathrm{root}, x, a)\pi_{x;a}\phi_x \left[= \sum_{x,a} \alpha_n(m, x, a)\beta_n(m, x, a)\right], \tag{S5}$$

- the posterior state probability at node $m$:

$$P^{(m)}(x, a \mid C_n, \theta) = \frac{\alpha_n(m, x, a) \cdot \beta_n(m, x, a)}{P(C_n \mid \theta)}, \tag{S6}$$

- the posterior joint state probability for both ends of branch $m$:

$$P^{(m)}(x, a, y, b \mid C_n, \theta) = \frac{\alpha_n(m, x, a)P(x, a|y, b, t_m R)\beta_n(m, y, b)}{P(C_n \mid \theta)}. \tag{S7}$$

##### 1.1.2 Scaling

In the inside-outside algorithm, the inside and outside variables can be extremely small, possibly resulting in underflow; therefore, actual calculations are often performed on a logarithmic scale, taking the logarithms of the Equations (S2) and (S4). However, this raises another issue of computational cost owing to the frequent call of log-sum-exp, which is generally slower than addition and multiplication.

Here, we describe an alternative approach using scaling factors, which involves dividing the variables by appropriate factors at each step in the recursion to ensure that they fall within a computationally tractable range. These factors can be rationally chosen based on the Bayes' theorem.

First, we precomputed the prior state probability at each tree node.

$$P^{(m)}(x, a \mid \theta) = \begin{cases} \pi_{x;a}\phi_x & (m \text{ is root}) \\ \sum_{y,b} P(x, a \mid y, b, t_m R) \cdot P^{(\mathrm{pa}(m))}(y, b) & (\text{otherwise}) \end{cases} \tag{S8}$$

The extra cost is some matrix-vector products independent of $N$, which is minimal. The transition probabilities are already required in the original inside-outside algorithm.

Subsequently, we recursively define the scaling factors $\kappa_{n,m}$ for each node as follows:

$$\kappa_{n,m} \cdot \hat{\alpha}_n(m, y, b) = \prod_{m' \in \mathrm{ch}(m)} \sum_{x,a} \hat{\alpha}_n(m', x, a) \cdot P(x, a \mid y, b, t_{m'} R), \tag{S9}$$

such that:

$$\sum_{x,a} P^{(m)}(x, a \mid \theta) \cdot \hat{\alpha}_n(m, x, a) = 1. \tag{S10}$$

We define the rescaled outside variables $\hat{\beta}_n$ using $\kappa_{n,m}$.

$$\hat{\beta}_n(m,x,a) = \frac{1}{\kappa_{n,\mathrm{pa}(m)}} \sum_{y,b} \Bigg[ P(x,a \,|\, y,b,t_m R) \cdot \hat{\beta}_n(\mathrm{pa}(m),y,b) \\ \times \prod_{m' \in \mathrm{sib}(m)} \sum_{x',a'} \hat{\alpha}_n(m',x',a') \cdot P(x',a' \,|\, y,b,t_{m'} R) \Bigg] \tag{S11}$$

Our scaling method provides a probabilistic interpretation. Equations (S9) implies $\alpha_n(m,x,a) = \left(\prod_{m' \in \mathrm{sub}(m)} \kappa_{n,m'}\right) \cdot \hat{\alpha}_n(m,x,a)$, where $\mathrm{sub}(m)$ is the set of nodes in the subtree rooted by $m$. Together with Equation (S10), we obtain

$$\begin{aligned} P(\mathrm{inside}_m(C_n) \,|\, \theta) &= \sum_{x,a} P^{(m)}(x,a \,|\, \theta) \alpha_n(m,x,a) \\ &= \left(\prod\nolimits_{m' \in \mathrm{sub}(m)} \kappa_{n,m'}\right) \underbrace{\sum_{x,a} P^{(m)}(x,a \,|\, \theta) \hat{\alpha}_n(m,x,a)}_{=1}. \end{aligned} \tag{S12}$$

It follows that from the definition of the inside variables (Equation (S1)),

$$\hat{\alpha}_n(m,x,a) = P^{(m)}(x,a \,|\, \mathrm{inside}_m(C_n), \theta). \tag{S13}$$

Also, Equation (S4) implies that $\beta_n(m,x,a) = \left(\prod_{m' \in \mathrm{out}(m)} \kappa_{n,m'}\right) \cdot \hat{\beta}_n(m,x,a)$, where $\mathrm{out}(m)$ is the set of tree nodes excluding $\mathrm{sub}(m)$. Therefore, from Equations (S1), (S3), and (S12), we obtain

$$\begin{aligned} \sum_{x,a} \hat{\alpha}_n(m,x,a) \hat{\beta}_n(m,x,a) &= \sum_{x,a} \alpha_n(m,x,a) \beta_n(m,x,a) \Big/ \prod_m \kappa_{n,m} \\ &= P(C_n \,|\, \theta) \,/\, P(\mathrm{inside}_{\mathrm{root}}(C_n) \,|\, \theta) = 1. \end{aligned} \tag{S14}$$

Due to Equations (S10) and (S14), $\hat{\alpha}_n$ and $\hat{\beta}_n$ are resistant to underflow.

#### 1.2 Details of EM algorithm

##### 1.2.1 E-step: on expectation of sufficient statistics

Here, we describe how to compute the expected value of the sufficient statistics in the $Q$-function of the main text. Since we obtained the posterior probability of the states at both ends of each branch (S7), we must solve is the behaviour of CTMC given the start and end points.

$$\begin{cases} N_\bullet^{(m)}(\boldsymbol{i},\boldsymbol{j} \,|\, C_n,\theta) = \sum_{\boldsymbol{a},\boldsymbol{b}} N_\pm^{(m)}(\boldsymbol{i},\boldsymbol{j} \,|\, \boldsymbol{a},\boldsymbol{b},\theta) P^{(m)}(\boldsymbol{a},\boldsymbol{b} \,|\, C_n,\theta) \\ F_{\mathrm{dur}}^{(m)}(\boldsymbol{i} \,|\, C_n,\theta) = \sum_{\boldsymbol{a},\boldsymbol{b}} F_{\mathrm{dur}}^{(m)}(\boldsymbol{i} \,|\, \boldsymbol{a},\boldsymbol{b},\theta) \quad P^{(m)}(\boldsymbol{a},\boldsymbol{b} \,|\, C_n,\theta) \\ n_{\mathrm{root}}(\boldsymbol{a} \,|\, C_n,\theta) = P(\boldsymbol{a} \,|\, C_n,\theta) \end{cases} \tag{S15}$$

Here, $N_\pm$ and $N_{\rm sw}$ are collectively denoted as $N_\bullet$ because their expressions are equivalent. To save space, bold letters $\boldsymbol{i}$, $\boldsymbol{j}$, $\boldsymbol{a}$ and $\boldsymbol{b}$ denote the compound states $(k,i)$, $(l,j)$, $(x,a)$ and $(y,b)$, respectively ($i=j$ for $N_\pm$ and $k=l$ for $N_{\rm sw}$); the superscript "old" is omitted from $\theta$, for simplicity.

The sufficient statistics on the right-hand side of the above equations have analytical forms (see, for example, Kiryu 2011 for a detailed derivation).

$$\begin{cases} N_\bullet^{(m)}(\boldsymbol{i},\boldsymbol{j}\,|\,\boldsymbol{a},\boldsymbol{b},\theta) = \dfrac{1}{[e^{t_m R}]_{\boldsymbol{a};\boldsymbol{b}}} \displaystyle\int_0^{t_m} \underbrace{[e^{(t_m-t)R}]_{\boldsymbol{a};\boldsymbol{i}}}_{\substack{\boldsymbol{a}\leftarrow\boldsymbol{i} \\ \text{in } [t,t_m]}} \underbrace{[e^{tR}]_{\boldsymbol{j};\boldsymbol{b}}}_{\substack{\boldsymbol{j}\leftarrow\boldsymbol{b} \\ \text{in } [0,t]}} \underbrace{R_{\boldsymbol{i};\boldsymbol{j}} dt}_{\substack{\boldsymbol{i}\leftarrow\boldsymbol{j} \\ \text{at } t}} \\ t_m F_{\rm dur}^{(m)}(\boldsymbol{i}\,|\,\boldsymbol{a},\boldsymbol{b},\theta) = \dfrac{1}{[e^{t_m R}]_{\boldsymbol{a};\boldsymbol{b}}} \displaystyle\int_0^{t_m} \underbrace{[e^{(t_m-t)R}]_{\boldsymbol{a};\boldsymbol{i}}}_{\substack{\boldsymbol{a}\leftarrow\boldsymbol{i} \\ \text{in } [t,t_m]}} \underbrace{[e^{tR}]_{\boldsymbol{i};\boldsymbol{b}}}_{\substack{\boldsymbol{i}\leftarrow\boldsymbol{b} \\ \text{in } [0,t]}} \underbrace{dt}_{\substack{\text{stay at } \boldsymbol{i} \\ \text{at } t}} \end{cases} \tag{S16}$$

One of the most efficient method to numerically solve these integrals is to use the eigendecomposition of $R = U\Lambda U^{-1}$, which simplifies the integrals to convolutions of univariate exponential functions. For a plain CTMC for ordinal quantities, where $R$ is a tridiagonal matrix with all sub-diagonal elements positive, eigendecomposition is always feasible in real space as $R$ can be converted to a real symmetric tridiagonal matrix with a similarity transformation; real symmetric tridiagonal matrices always have eigendecomposition in real space. However, in the Markov-modulated CTMC case, we cannot guarantee that eigendecomposition is always possible. The (counter) proof of this will be the subject of future research.

Other approaches are based on the Frechét derivative of the matrix exponential. Rather, the above integrals are the integral representations of the (partial) Frechét derivative itself. More specifically, let $A$ be a square matrix and $E(i,j)$ be a matrix of the same size as $A$ where the $(i,j)$-th element is 1 and all others are 0. Then, the partial Frechét derivative in the direction of $E(i,j)$ is given as:

$$\frac{\partial}{\partial A_{i,j}} \exp(tA) = \int_0^t \exp((t-t')A)E(i,j)\exp(t'A)dt'. \tag{S17}$$

Several methods have been proposed to solve this derivative, including the block enlargement and the Scaling-Padé-Squaring (Al-Mohy and Higham, 2009).

Herein, we selected the eigendecomposition method primarily due to its computational speed and switched to the block enlargement method in cases of numerical instability. Only the former method was used to measure the execution times.

##### 1.2.2 M-step: on maximisation of $Q$-function

In the M-step, we maximise the $Q$-function in the main text subject to the following constraints:

$$\begin{cases} \displaystyle\sum_i \pi_{x;a} - 1 =: G_{\pi;x}(\pi_x) = 0 \quad (x = 1,\cdots,K) \\ \displaystyle\sum_x \phi_x \ - 1 =: G_\phi(\phi) \quad = 0 \end{cases} \tag{S18}$$

Using the method of Lagrange multipliers, we obtain the necessary conditions for the (local) maxima.

$$
\begin{cases}
\dfrac{\partial Q}{\partial \pi_{k;i}} - \lambda_{\pi;x} \dfrac{\partial G_{\pi;x}}{\partial \pi_{x,a}} = \sum_{n} n_{\mathrm{root}}\big(x, a \,\big|\, C_n, \theta^{\mathrm{old}}\big) - \lambda_{\pi;x} = 0 \\
\dfrac{\partial Q}{\partial \phi_x} - \lambda_{\phi} \dfrac{\partial G_{\phi}}{\partial \phi_x} = \sum_{n,a} n_{\mathrm{root}}\big(x, a \,\big|\, C_n, \theta^{\mathrm{old}}\big) - \lambda_{\phi} = 0 \\
\dfrac{\partial Q}{\partial r_{k;i,j}} = \dfrac{1}{r_{k;i,j}} \sum_{n,m} N_{\pm}^{(m)}\big(k, i, j \,\big|\, C_n, \theta^{\mathrm{old}}\big) - \sum_{n,m} t_m F_{\mathrm{dur}}^{(m)}\big(k, j \,\big|\, C_n, \theta^{\mathrm{old}}\big) = 0 \\
\dfrac{\partial Q}{\partial s_{k,l}} = \dfrac{1}{s_{k,l}} \sum_{n,m,j} N_{\mathrm{sw}}^{(m)}\big(k, l, j \,\big|\, C_n, \theta^{\mathrm{old}}\big) - \sum_{n,m,j} t_m F_{\mathrm{dur}}^{(m)}\big(l, j \,\big|\, C_n, \theta^{\mathrm{old}}\big) = 0,
\end{cases}
\tag{S19}
$$

where $\lambda_{\pi;k}$ and $\lambda_{\phi}$ are Lagrange multipliers. Recall that $[\operatorname{diag} R]_{l,j} = -\sum_{i \neq j} r_{l;i,j} - \sum_{k \neq l} s_{k,l}$ for all $(l, j)$.

By solving this system (S19) subject to constraints (S18), we obtain the intuitive update formula as follows:

$$
\begin{aligned}
\pi_{x;a} &= \frac{\sum_n n_{\mathrm{root}}\big(x, a \,\big|\, C_n, \theta^{\mathrm{old}}\big)}{\sum_{n,a'} n_{\mathrm{root}}\big(x, a' \,\big|\, C_n, \theta^{\mathrm{old}}\big)}, \quad
& r_{k;i,j} &= \frac{\sum_{n,m} N_{\pm}^{(m)}\big(k, i, j \,\big|\, C_n, \theta^{\mathrm{old}}\big)}{\sum_{n,m} t_m F_{\mathrm{dur}}^{(m)}\big(k, j \,\big|\, C_n, \theta^{\mathrm{old}}\big)}, \\
\phi_x &= \frac{\sum_{n,a} n_{\mathrm{root}}\big(x, a \,\big|\, C_n, \theta^{\mathrm{old}}\big)}{\sum_{n,k',i} n_{\mathrm{root}}\big(k', i \,\big|\, C_n, \theta^{\mathrm{old}}\big)}, \quad
& s_{k,l} &= \frac{\sum_{n,m,i} N_{\mathrm{sw}}^{(m)}\big(k, l, i \,\big|\, C_n, \theta^{\mathrm{old}}\big)}{\sum_{n,m,j} t_m F_{\mathrm{dur}}^{(m)}\big(l, j \,\big|\, C_n, \theta^{\mathrm{old}}\big)}.
\end{aligned}
\tag{S20}
$$

#### 1.3 Practical notes

##### 1.3.1 Initial value selection

The EM algorithm has an initial value dependency. This algorithm ensures the convergence of the log-likelihood; however, in general, this does not necessarily mean convergence to the global optimum. In other words, starting from different initial values, the parameters may converge to different stationary points (local optima or even saddle points).

Accordingly, we recommend using multiple initial values in our model. In this study, we adopted the emEM method (Biernacki *et al.*, 2003), in which multiple short EM runs were executed, and the best run in terms of likelihood was selected as the initial value. We prepared 10 initial values for each condition, with the emEM initialisation configured to perform 25 short runs for each.

##### 1.3.2 Acceleration of EM algorithm

To reduce the number of likelihood evaluations, we integrated a series acceleration method into the EM algorithm. Specifically, we adopt the parabolic acceleration of the EM algorithm (P-EM; Berlinet and Roland, 2012). In brief, the P-EM algorithm seeks to update parameters in bigger strides by extrapolating the trajectory of parameters with the quadratic Bézier curve (i.e., a parabola).

1. Draw the quadratic Bézier curve using parameters in the last three updates $\theta^{(t)}$, $\theta^{(t-1)}$ and $\theta^{(t-2)}$:

$$\theta_B(\tau) = (1-\tau)^2\theta^{(t-2)} + 2\tau(1-\tau)\theta^{(t-1)} + \tau^2\theta^{(t)}$$

2. Perform a grid search on the curve as long as a better likelihood is yielded. This time, we used a geometric grid $\tau = 1, 1+ha, 1+ha^2, \cdots$ ($h = 0.1$ and $a = 1.5$ as recommended in Berlinet and Roland (2012)).

$$\hat{\tau} = \min\{\tau_i \mid \log L(\theta_B(\tau_i)) \geq \log L(\theta_B(\tau_{i+1}))\}$$

3. Update parameters twice following the normal procedure of the EM algorithm to stabilise trajectory:

$$\theta^{(t+1)} = \mathrm{EM}(\mathrm{EM}(\theta_B(\hat{\tau})))$$

4. Set $t$ to $t+1$ and go back to Step 1 unless convergence.

In Step 2, we heuristically skipped the likelihood computation for some $\tau_i$ ($i > 0$) as long as $\|\theta^{(t)} - \theta_B(\tau_i)\|_2$ was less than the last update in Step 3 of the previous cycle. The rationale for this heuristic is that Step 2 is intended to increase the step size of the updates over normal EM updates.

The monotonic increase in likelihood still holds for the P-EM algorithm and our heuristics, which ensures the convergence of these algorithms. However, there is no theoretical guarantee of whether they will converge to the same values as the plain EM algorithm.

In this study, we used the modified P-EM algorithm only in the main long run while adopting plain EM for emEM initialisation.

##### 1.3.3 Stopping criteria

During the analyses in this study, we determined the convergence of the EM algorithm(s) based on the step size of parameter updates.

In the emEM initialisation, we stopped the short runs of the plain EM algorithm if (1) it reached 100 iterations or (2) all the parameters were close to the previous step with a relative tolerance of 0.001. In the main model fitting, we stopped the modified P-EM algorithm in Step 3 if (1) the likelihood evaluation exceeded 5,000 times or (2) all parameters were close to the previous step with a relative tolerance of $10^{-6}$ or an absolute tolerance of $10^{-6}$. The likelihood evaluations for the rejected parameters in acceleration Step 2 were also included in the count.

#### 1.4 Parameter regularisation and MAP estimation

To improve the stability of the parameter fitting, we derived an EM algorithm for maximum a posteriori (MAP) estimation. In the MAP estimation, the posterior probability of the parameters was optimised $P(\theta \mid C) \propto P(C \mid \theta)P(\theta)$ instead of the data likelihood $P(C \mid \theta)$ in the ML estimation. To perform MAP estimation using the EM algorithm, we modified the $Q$-function of the ML estimation by adding a term $P(\theta)$.

For computational convenience, we assumed the following prior distributions:

$$\begin{aligned} \boldsymbol{\pi}_k &\sim \mathrm{Dirichlet}(\boldsymbol{\alpha}), \quad & r_{k;i,j} &\sim \Gamma(k_\pm, \lambda_\pm), \\ \boldsymbol{\phi} &\sim \mathrm{Dirichlet}(\boldsymbol{\beta}), \quad & s_{k,l} &\sim \Gamma(k_{\mathrm{sw}}, \lambda_{\mathrm{sw}}). \end{aligned} \tag{S21}$$

Then, the M-step in MAP estimation falls into a "smooth" version of the M-step in ML estimation (S20):

$$\begin{aligned} \pi_{k;i} &= \frac{\{n_{\mathrm{root}}(k, i \mid \cdots)\} + \alpha_i}{\{n_{\mathrm{root}}(k, \cdot \mid \cdots)\} + \sum_{i'} \alpha_{i'}}, \quad & r_{k;i,j} &= \frac{\{N_\pm(k, i, j \mid \cdots)\} + k_\pm - 1}{\{tF_{\mathrm{dur}}(k, j \mid \cdots)\} + 1/\lambda_\pm}, \\ \phi_k &= \frac{\{n_{\mathrm{root}}(k, \cdot \mid \cdots)\} + \beta_k}{\{n_{\mathrm{root}}(\cdot, \cdot \mid \cdots)\} + \sum_{k'} \beta_{k'}}, \quad & s_{k,l} &= \frac{\{N_{\mathrm{sw}}(k, l, \cdot \mid \cdots)\} + k_{\mathrm{sw}} - 1}{\{tF_{\mathrm{dur}}(l, \cdot \mid \cdots)\} + 1/\lambda_{\mathrm{sw}}}, \end{aligned} \tag{S22}$$

where "$\{\cdots\}$" indicates the abbreviation of the corresponding parts in Equation (S20).

MAP estimation boils down to ML estimation at the limits of $k_\pm, k_{\mathrm{sw}} \to 1, \lambda_\pm, \lambda_{\mathrm{sw}} \to \infty, \boldsymbol{\alpha} \to \mathbf{1}, \boldsymbol{\beta} \to \mathbf{1}$. Moreover, when the shape parameter $k$ is 1, the gamma distribution of the rate parameters is reduced to an exponential distribution, which implies sparse modelling. This may help to mitigate the complexity of the model, particularly for switching rates consisting of $O(K^2)$ parameters.

In practice, we configured $\boldsymbol{\alpha} = [1.1, \cdots, 1.1]$, $\boldsymbol{\beta} = [1.1, \cdots, 1.1]$, $(k_\pm, \lambda_\pm) = (2, \overline{\Delta}_{\mathrm{MP}} / \sum t_m / 2)$ and $(k_{\mathrm{sw}}, \lambda_{\mathrm{sw}}) = (3, \overline{\Delta}_{\mathrm{MP}} / \sum t_m / 3)$, where $\overline{\Delta}_{\mathrm{MP}}$ is the number of parsimonious changes in copy number averaged over all gene families.

### 2 Supplementary Notes

#### 2.1 Related works

The idea of Markov modulation has been applied to the models of molecular sequence evolution (Tuffley and Steel, 1998; Gascuel and Guindon, 2007; Baele *et al.*, 2021) and other phylogenetic comparative methods (Beaulieu *et al.*, 2013; Boyko and Beaulieu, 2021; Höhna *et al.*, 2019).

Regarding sequences, covarion-style models (Tuffley and Steel, 1998; Galtier, 2001) can be regarded as special cases of Markov modulation, where there are two categories, with one representing the invariant mode. More general cases are described in Gascuel and Guindon (2007) and Baele *et al.* (2021). In these models, both substitution and switching processes were assumed to be in a steady state. In contrast, our model was not assumed as such since gene content evolution may be nonstationary.

In the context of the evolution of discrete characters, the hidden rates model (Beaulieu *et al.*, 2013) and its generalisation (Boyko and Beaulieu, 2021), which are implemented in the R package corHMM, make the most of Markov modulation to account for rate heterogeneity. Recently, corHMM was applied to infer the correlated evolution of gene pairs Diao *et al.* (2024). The major difference between corHMM and our model is how the model is applied to the data, owing to the difference in the scope of analysis. The authors focused on analysing an individual character or a combination of characters. In contrast, we handled all gene families collectively as independent samples from an identical model to extract the overall evolutionary patterns. The birth-death-shift model of species diversification by Höhna *et al.* (2019) is also analogous in that it relies on unobservable hidden states to enable a shift in the rates of the birth-death process.

Our model is reduced to a mixture model (Mirage; Fukunaga and Iwasaki, 2021) at the limit of $s_{k,l} \to 0$. In other words, our model is a natural generalisation of Mirage. At the limit of $s_{k,l} \to \infty$, on the other hand, we qualitatively expect that our model simplifies to a plain CTMC with a single rate category. Similar arguments have been made in the literature (Galtier, 2001; Gascuel and Guindon, 2007) but without mathematical proof.

### 3 Supplementary Results

#### 3.1 Estimation of simulated datasets

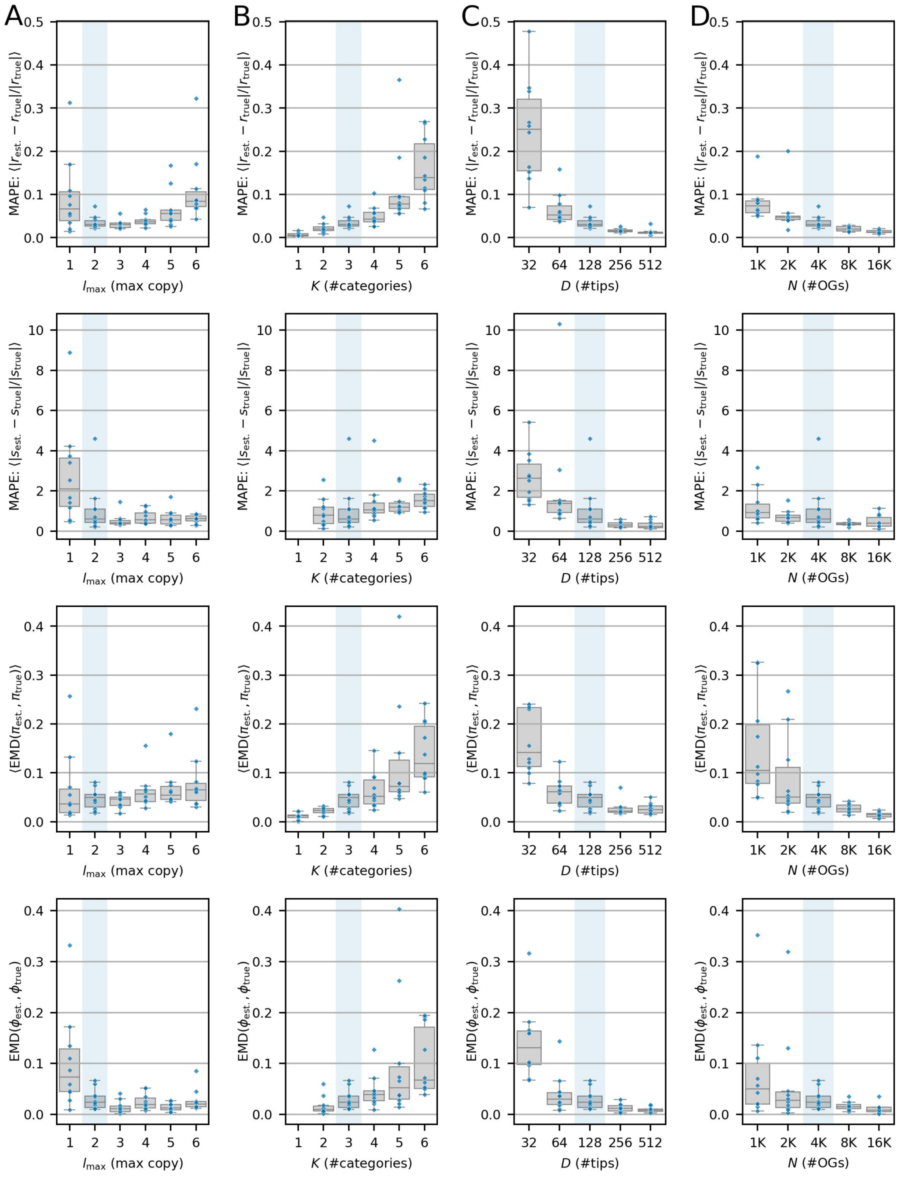


**Supplementary Fig. S1.** Distributions of the errors in parameter estimation for the simulated datasets. The y-axes represent the mean absolute percentage error (MAPE) for rate parameters and the (mean) Earth mover's distance (EMD) for root probabilities $\pi$ and $\phi$. Each data point corresponds to the estimation of an individual dataset. The box shows the quartiles, and the whiskers indicate the range of data excluding outliers (data >1.5 inter-quartile range away from the box). All conditions highlighted in blue in each panel are identical.

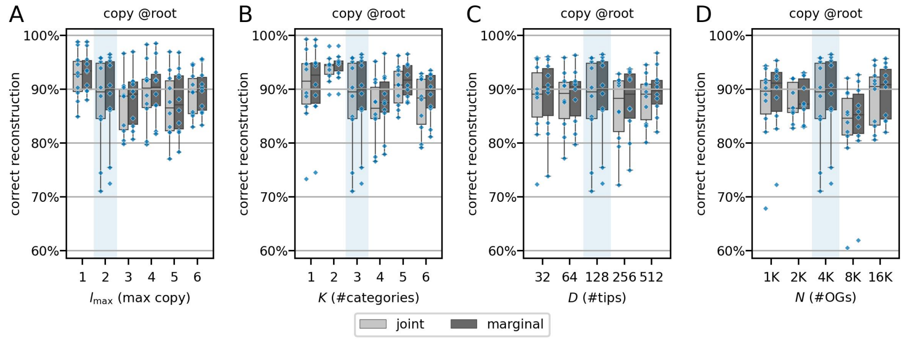


**Supplementary Fig. S2.** Distribution of the percentage of gene families for which copy number was correctly estimated at the tree root of each simulated dataset.

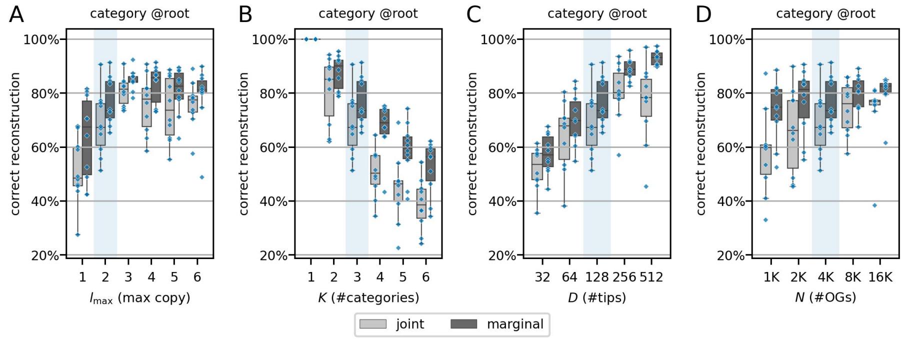


**Supplementary Fig. S3.** Distribution of the percentage of gene families for which rate category was correctly estimated at the tree root of each simulated dataset.

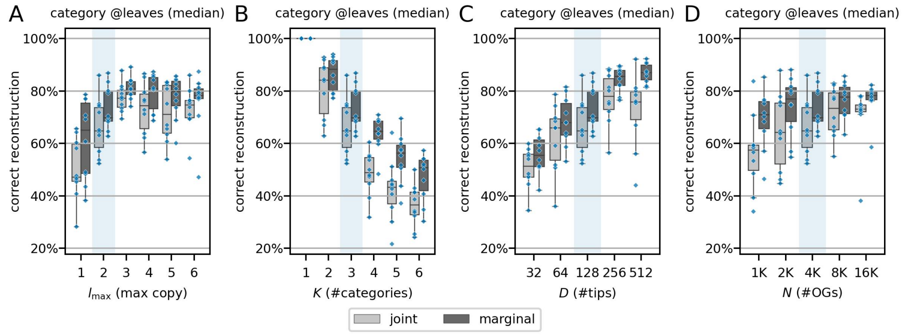


**Supplementary Fig. S4.** Distribution of the median percentage of gene families for which rate category was correctly estimated at the tree leaves of each simulated dataset.

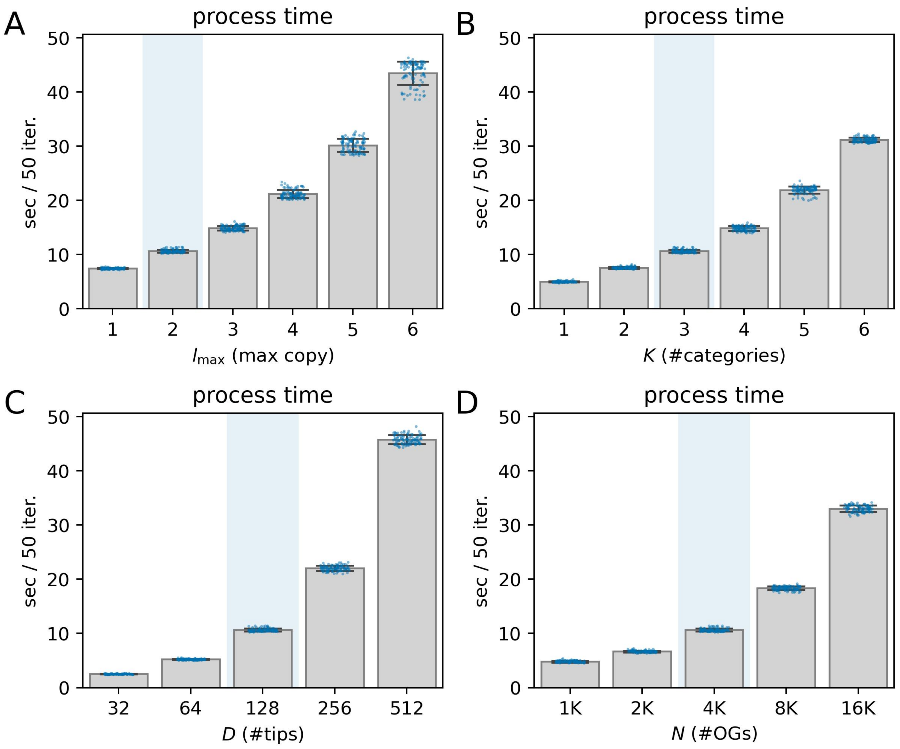


**Supplementary Fig. S5.** Execution time of the proposed model measured using simulation data. The y-axis represents the time (seconds) required for 50 EM iterations, while the x-axis shows (A) $l_{\max}$, (B) $K$, (C) $D$ and (D) $N$. Swarm plots depict results from 10 datasets $\times$ 10 replicates for each condition, with bar plots indicating the mean $\pm$ SD. The conditions highlighted in blue in each panel are the identical baseline conditions.

#### 3.2 Summary of the empirical datasets

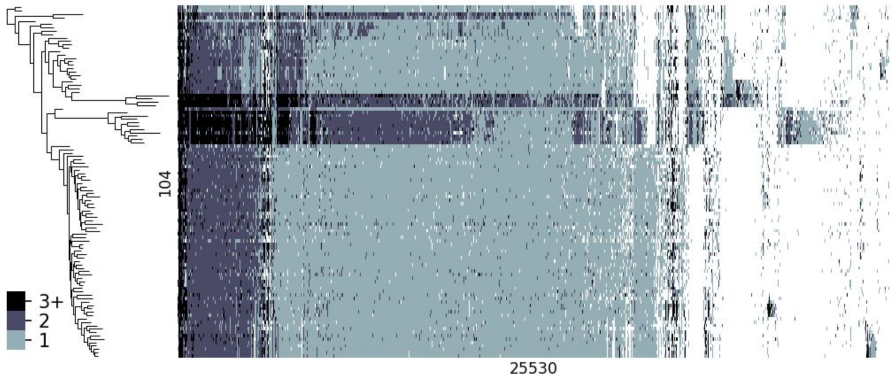


**Supplementary Fig. S6.** The overview of the ray-finned fish dataset. The copy-number profile (H: species × W: gene families) is shown as a heatmap, limiting $l_{\max}$ to 3.

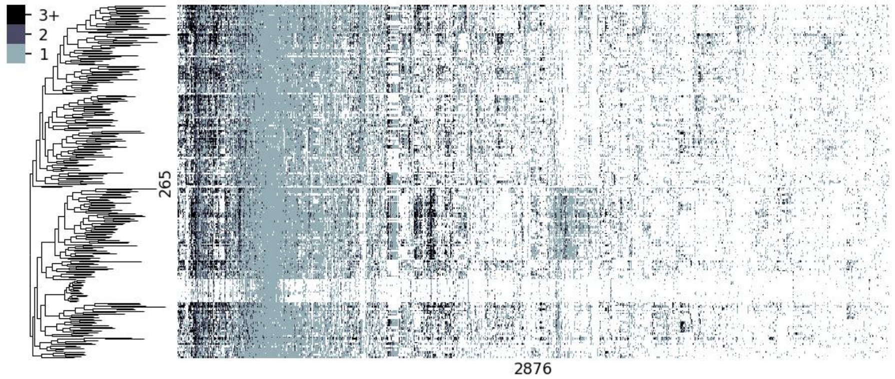


**Supplementary Fig. S7.** The overview of the bacteria domain-wide dataset. The copy-number profile (H: species × W: gene families) is shown as a heatmap, limiting $l_{\max}$ to 3.

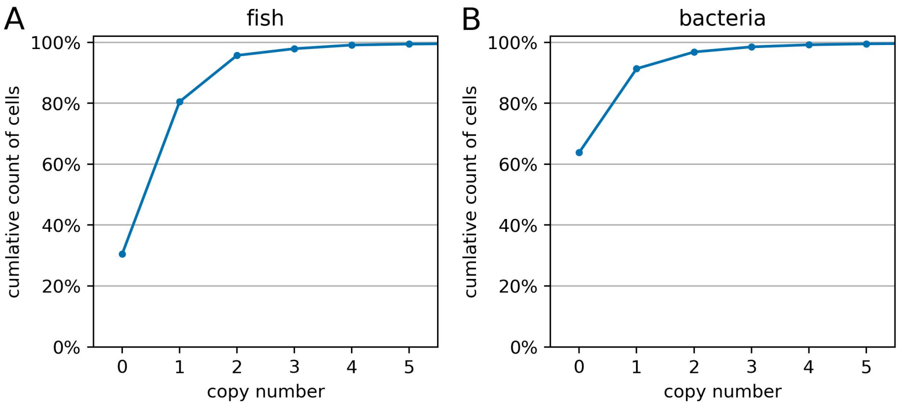


**Supplementary Fig. S8.** Cumulative distribution curve of the gene copy numbers in the empirical datasets. The x-axis represents gene copy numbers, and the y-axis indicates the percentage of the cells in ortholog tables whose value does not exceed the threshold.

#### 3.3 MAP estimation of empirical datasets

As shown in Supplementary Figure S9, the scores in the MAP estimation were almost tied to those in the ML estimation, except for the free-rate branch model. This means that modest prior distributions, as specified, have little effect on the goodness of fit. Thus, it is beneficial to perform MAP estimation if the estimation of our model is unstable. However, in general, overspecification of the prior distribution often leads to biased estimates.

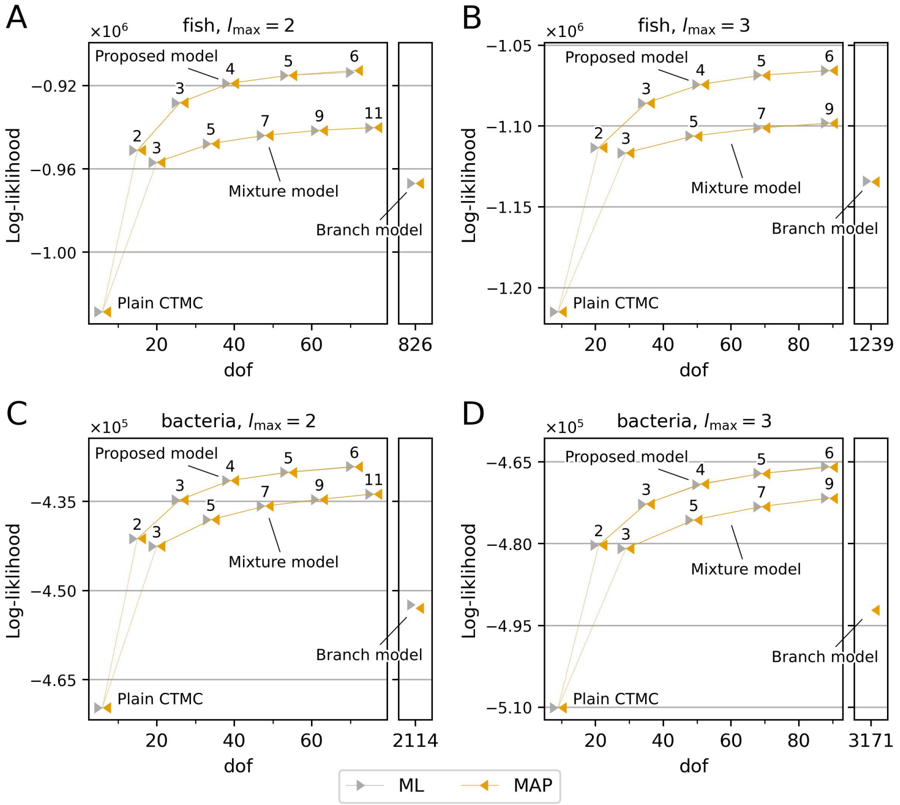


**Supplementary Fig. S9.** Comparison of model fit between ML and MAP estimations. The layout is the same as in Figure 2. The grey markers show the cross-validation scores for the ML estimation, reiterating Figure 2 for comparison. The orange markers show the scores for the MAP estimation.

#### 3.4 Ancestral state estimation

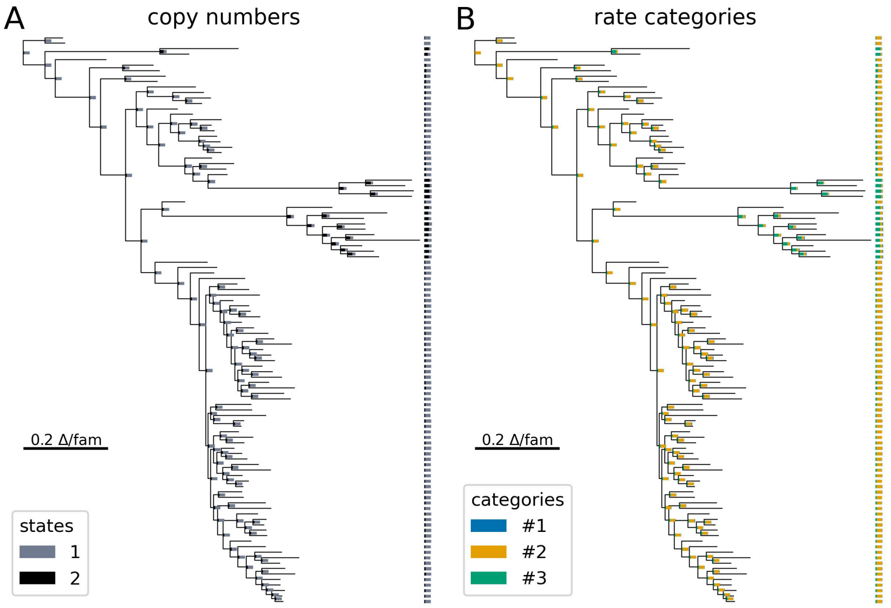


**Supplementary Fig. S10.** Ancestral state estimation of the fish dataset with marginal estimation. $l_{\max}$ and $K$ were set to 2 and 3, respectively. The tree is a phylogram whose branch lengths represent the changes in gene copy numbers per gene family approximated by the maximum parsimony. Markers attached to the internal and leaf nodes show the estimated composition of (A) gene copy numbers and (B) rate categories. The fractions for gene families estimated to be absent in each node have been excluded.

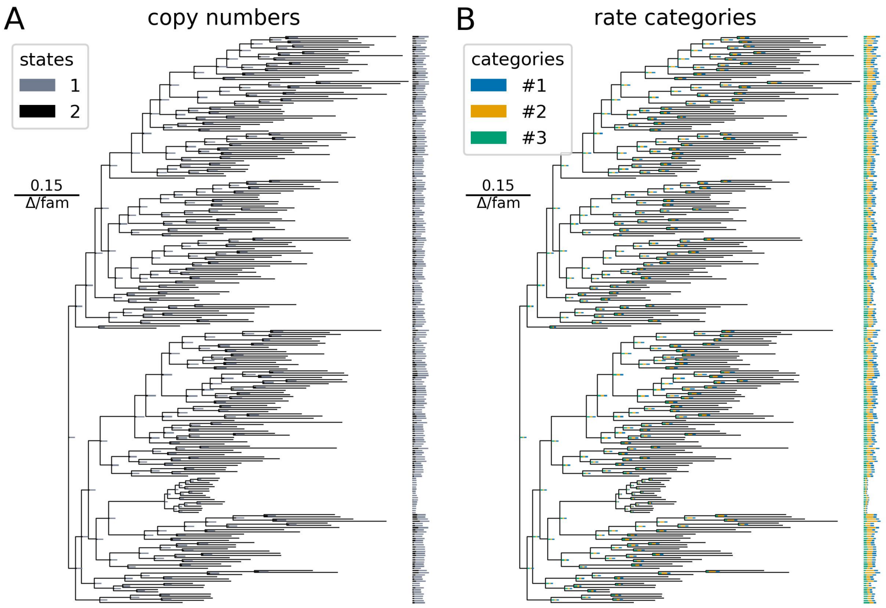


**Supplementary Fig. S11.** Ancestral state estimation of the bacteria dataset with marginal reconstruction, presented in the same way as Supplementary Figure S10.

#### 3.5 Branch lengths heavily affect estimation

We assumed that the branch length setting was likely to have a strong effect on model fit and ancestral state estimation. Recently, the importance of selecting branch-length sets in estimating ancestral states has been recognised (Wilson *et al.*, 2022), and this notion inevitably applies to our task. In principle, our model can handle rate heterogeneity among branches owing to Markov modulation; however, if the extent is excessive, it becomes difficult to do so with a limited number of rate categories. Therefore, we expected that it would be effective for ancestral state estimation to rescale branch lengths to equalise the scale of evolutionary rates, allowing rate categories to focus on more global and essential evolutionary patterns.

To approximate the extent of the evolutionary changes, we relied on maximum parsimony (MP). We calibrated the branch lengths by averaging the number of copy number changes per gene family over all possible MP reconstructions. These averages can be efficiently computed using a combinatorics version of the inside-outside algorithm, in which the (unweighted) number of possible reconstructions is calculated, instead of their probabilities. Specifically, this can be achieved by replacing the transition probability matrix in Equations (S2) and (S4) with a binary matrix that indicates the transitions of minimal cost.

Supplementary Figures S12 and S13 show the results of model fitting and ancestral state estimation for the fish dataset without branch-length adjustment. We followed the same procedure as in the main results (Figure 3; Supplementary Figure S10), except for using a chronogram as input rather than a phylogram with its branch lengths adjusted to the extent of changes in gene content. Compared with the main results, the performance of our model deteriorated significantly. The estimated parameters did not contain rate categories associated with whole-genome duplication (Supplementary Figure S12). Notably, in the near-root deep internal nodes, almost all gene families were estimated to have multiple copies (Supplementary Figure S13A), which was unlikely as this dataset was based on the Actinopterygii-level orthologous groups in OrthoDB. The estimated distribution of the rate categories was more homogenised throughout the entire tree (Supplementary Figure S13B) than when branch lengths were adjusted (Supplementary Figure S10B). These results demonstrate the importance of branch-length adjustments.

The correction method shown here is only one possibility; there is room to consider other methods in the future. MP underestimates evolutionary changes when dealing with distant evolutionary relationships as MP does not consider reversion and parallelism. Alternatively, optimising branch lengths jointly with the model parameters in the ML estimation may be possible. In the plain CTMC with a single rate category, it is known that optimal branch lengths represent the expected number of evolutionary changes (gene gain/loss) as a natural stochastic extension of the mean MP (Kiryu, 2011). Nevertheless, we could not adopt this approach herein since the ML estimation of branch lengths in our case also accounted for category switching, which may contain non-negligible errors (Supplementary Figures S1–S4). We argue that such issues concerning branch length should continue to be considered in the future developments of this methodology.

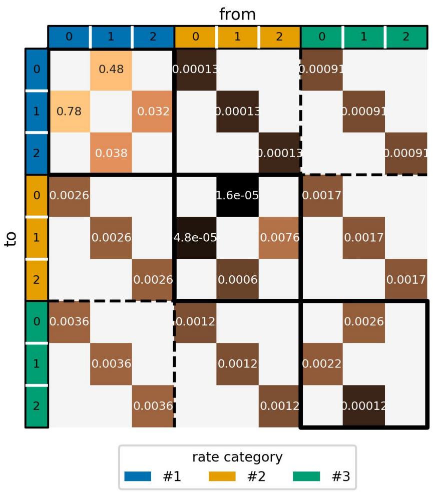


**Supplementary Fig. S12.** Transition rate matrix $R$ estimated for the fish dataset without adjustment of branch lengths.

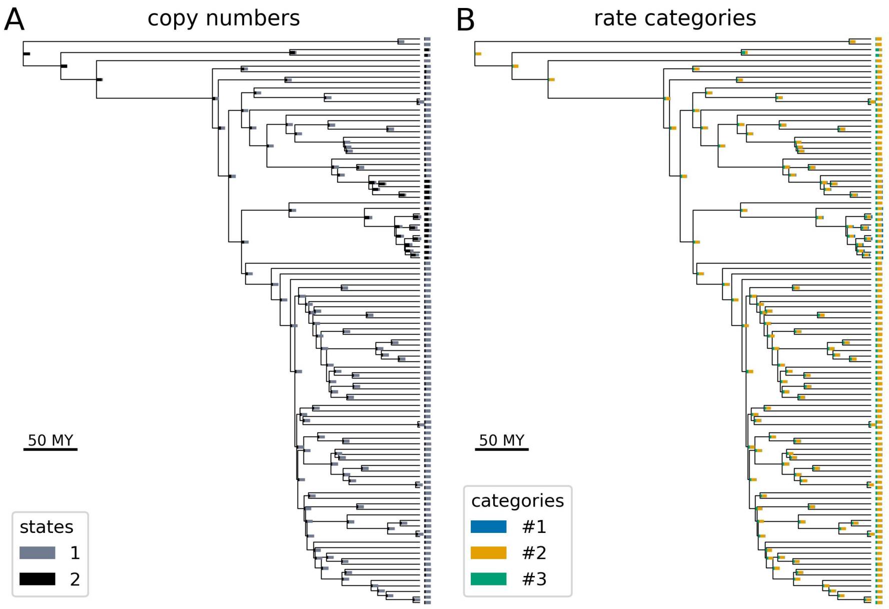


**Supplementary Fig. S13.** Ancestral state estimation of the fish dataset without adjustment of branch lengths. Shown in the same way as the previous figures, except that the tree is a chronograph, corresponding to the difference in preprocessing of the dataset.

#### 3.6 Acceleration by the parabolic EM algorithm

To evaluate the performance of the parabolic EM (P-EM) algorithm, we executed three EM algorithms (plain EM, original P-EM, and modified P-EM) using the same initial values and compared their fitting processes. We collected 10 runs for each method using the bacteria dataset, configuring our model as $l_{\mathrm{max}} = 2$ and $K = 3$. The initialisation and stopping criteria were the same as those adopted during the model fitting described in the main text.

Supplementary Figure S14 presents an overview of the fitting processes of the EM algorithms. The likelihood increased faster in the parabolic EM algorithms than in the plain EM algorithm (A). Our heuristics rendered the process more efficient in many cases by omitting unnecessary likelihood evaluations (A). As a result, the original P-EM algorithm reduced the number of likelihood evaluations required for convergence by 26% on average compared with the plain EM algorithm ($p = 0.002$; Wilcoxon signed-rank test), whereas the modified P-EM further reduced it by 35% compared with the original P-EM ($p = 0.027$) (B).

We investigated the trajectories of the parameters in the EM algorithms. Considering the label switching of rate categories, the parameters were dimensionally compressed using principal component analysis for visual inspection. At least in this experiment, all series converged to virtually the same values (C), suggesting that the P-EM algorithm and our modification did not affect the results.

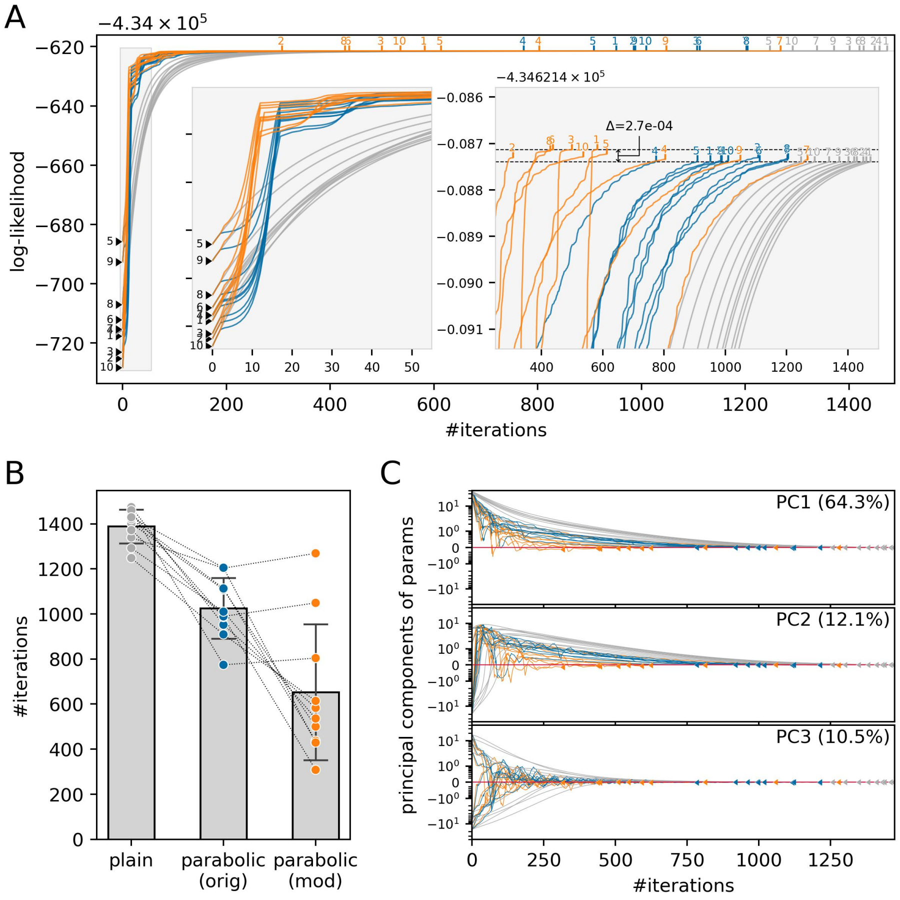


**Supplementary Fig. S14.** Summary of fitting processes in the EM algorithms. (A) Trajectories of the likelihood in each algorithm. The x- and y-axis indicate the number of likelihood evaluations and the log-likelihood at each time, respectively. Plots in grey, blue, and orange represent the plain EM, original P-EM, and modified P-EM algorithm, respectively. Rejected evaluations due to a decline in likelihood are not shown in plots, although they are included in the alignment of x-axis. Markers with a number indicate the group ID of the initial values. Inset panels illustrate the start and end of fitting in a magnified view. (B) The number of likelihood evaluations required for convergence in each algorithm. Each data point represents an individual EM run and runs from the same initial values are connected by a dotted line. Bar plots and error bars summarise the data in terms of mean ± SD. (C) Trajectories of parameters dimensionally compressed with the principal component analysis. The x-axis is the same as (A), and the y-axes are the first to third principal components of parameters. Markers show the convergence point of each run. The y-coordinate values are re-centred at the average convergence value of the plain EM runs, shown on an asinh scale.
